# Novel spirocyclic dimer, SpiD3, targets chronic lymphocytic leukemia survival pathways with potent preclinical effects

**DOI:** 10.1101/2024.01.31.578283

**Authors:** Alexandria P Eiken, Audrey L Smith, Sydney A Skupa, Elizabeth Schmitz, Sandeep Rana, Sarbjit Singh, Siddhartha Kumar, Jayapal Reddy Mallareddy, Aguirre A de Cubas, Akshay Krishna, Achyuth Kalluchi, M Jordan Rowley, Christopher R D’Angelo, Matthew A Lunning, R Gregory Bociek, Julie M Vose, Amarnath Natarajan, Dalia El-Gamal

## Abstract

Chronic lymphocytic leukemia (CLL) cell survival and growth is fueled by the induction of B-cell receptor (BCR) signaling within the tumor microenvironment (TME) driving activation of NF- κB signaling and the unfolded protein response (UPR). Malignant cells have higher basal levels of UPR posing a unique therapeutic window to combat CLL cell growth using pharmacological agents that induce accumulation of misfolded proteins. Frontline CLL therapeutics that directly target BCR signaling such as Bruton-tyrosine kinase (BTK) inhibitors (e.g., ibrutinib) have enhanced patient survival. However, resistance mechanisms wherein tumor cells bypass BTK inhibition through acquired BTK mutations, and/or activation of alternative survival mechanisms have rendered ibrutinib ineffective, imposing the need for novel therapeutics. We evaluated SpiD3, a novel spirocyclic dimer, in CLL cell lines, patient-derived CLL samples, ibrutinib-resistant CLL cells, and in the Eµ-TCL1 mouse model. Our integrated multi-omics and functional analyses revealed BCR signaling, NF-κB signaling, and endoplasmic reticulum stress among the top pathways modulated by SpiD3. This was accompanied by marked upregulation of the UPR and inhibition of global protein synthesis in CLL cell lines and patient-derived CLL cells. In ibrutinib-resistant CLL cells, SpiD3 retained its anti-leukemic effects, mirrored in reduced activation of key proliferative pathways (e.g., PRAS, ERK, MYC). Translationally, we observed reduced tumor burden in SpiD3-treated Eµ-TCL1 mice. Our findings reveal that SpiD3 exploits critical vulnerabilities in CLL cells including NF-κB signaling and the UPR, culminating in profound anti-tumor properties independent of TME stimuli.

**STATEMENT OF SIGNIFICANCE:** SpiD3 demonstrates cytotoxicity in CLL partially through inhibition of NF-κB signaling independent of tumor-supportive stimuli. By inducing the accumulation of unfolded proteins, SpiD3 activates the UPR and hinders protein synthesis in CLL cells. Overall, SpiD3 exploits critical CLL vulnerabilities (i.e., the NF-κB pathway and UPR) highlighting its use in drug-resistant CLL.

**VISUAL ABSTRACT:** 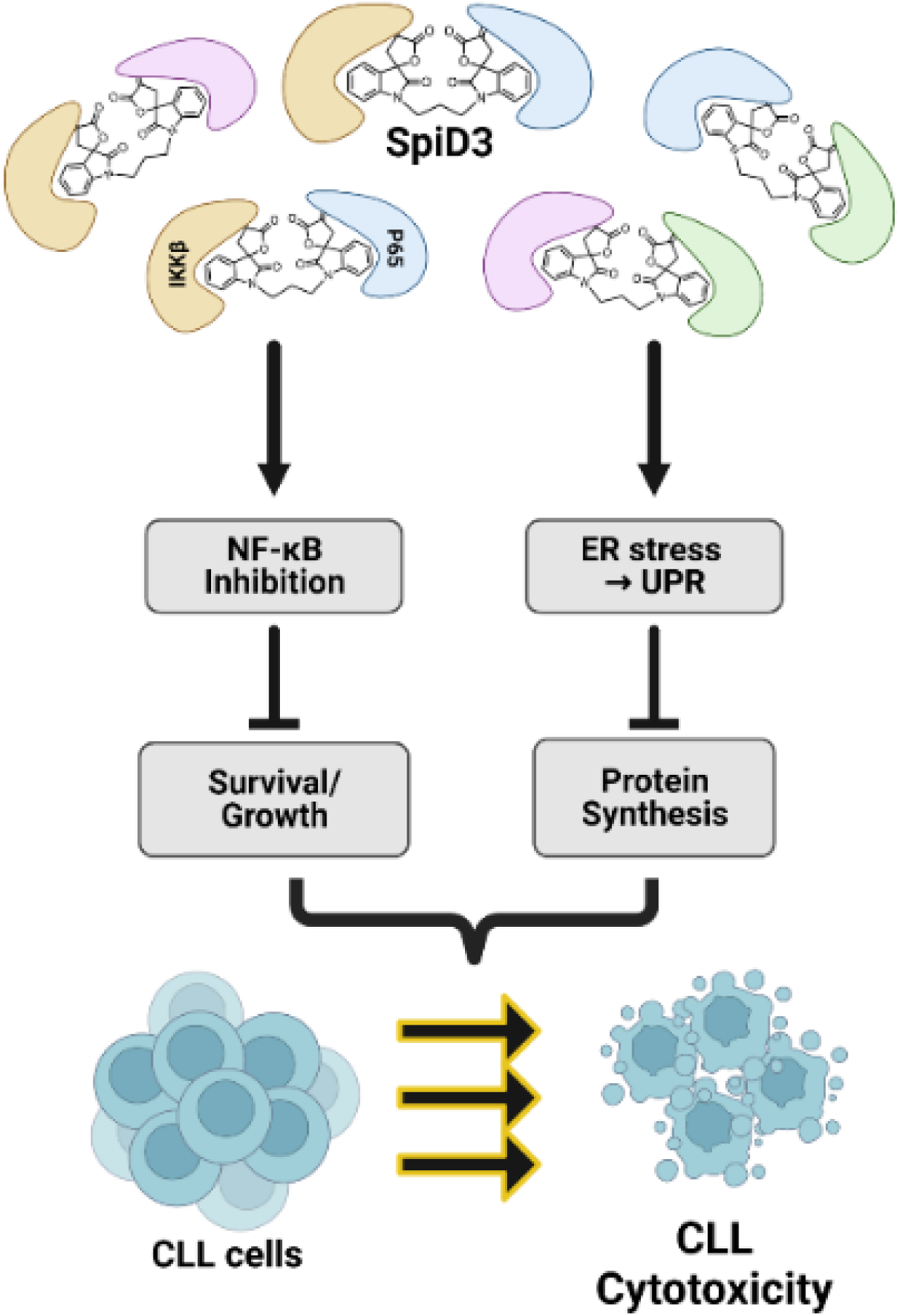

## INTRODUCTION

Chronic lymphocytic leukemia (CLL) is an incurable disease characterized by the accumulation of mature CD5^+^ B-cells in peripheral blood, bone marrow, and secondary lymphoid tissues (e.g., spleen and lymph nodes) [1]. The tumor microenvironment (TME) provides a protective niche supporting CLL cell survival and proliferation through activation of key pathways including B-cell receptor (BCR) and toll-like receptor (TLR) signaling [2, 3]. Antigenic stimulation activates BCR signaling which induces MYC activity and protein translation in CLL cells [4, 5]. Furthermore, crosstalk with surrounding TME cells, such as T-cells [2] and monocyte-derived nurse-like cells [6] create a permissive milieu protecting CLL cells from drug-induced apoptosis. These pro-survival/proliferation signals converge on the constitutively active nuclear factor kappa-light-chain-enhancer of activated B cells (NF-κB) pathway [2, 3]. Small molecule inhibitors targeting BCR signaling such as Bruton-tyrosine kinase (BTK) inhibitors (e.g., ibrutinib) have become the forefront of CLL therapeutics in the last decade. Despite the unprecedented clinical success of ibrutinib, resistance mechanisms wherein tumor cells bypass BTK inhibition through acquired BTK mutations (e.g., C481S) and/or activation of alternative survival mechanisms (e.g., PRAS/AKT/mTOR, ERK1/2, NF-κB), have rendered ibrutinib ineffective for many patients, imposing the need for novel therapeutics [7].

When the load of unfolded proteins is greater than the endoplasmic reticulum’s (ER) capacity to handle, the unfolded protein response (UPR) is activated [8, 9]. The UPR is a multifunctional response pathway that senses and adapts to ER stress partially through the eukaryotic translation initiation factor 2-alpha kinase 3 (PERK) and inositol-requiring enzyme 1α (IRE1α) pathways [8, 9]. PERK activation results in eIF2α phosphorylation, thereby inhibiting mRNA translation, reducing the load of newly synthesized proteins [8, 9]. Activated IRE1α splices XBP1 mRNA, which encodes a functional transcription factor that regulates genes involved in maintaining ER balance [8]. Cancer cells exhibit higher basal levels of unfolded proteins rendering them more vulnerable to UPR activation-induced apoptosis [9, 10]. Several studies have shown that pharmacological inducers of the UPR promote CLL apoptosis in vitro [11, 12].

Our recent studies identified spirocyclic dimers (SpiDs) containing α-methylene-γ-butyrolactone functionality as novel anticancer agents [10, 13, 14]. SpiDs covalently modify NF-κB proteins, P65 and IKKβ, by targeting unique surface-exposed cysteine (SEC) residues. Owing to its dimer structure, SpiD7 (7-carbon-linker SpiD) binds to SEC residues on cellular proteins to mimic misfolded proteins thereby activating the UPR and selectively inducing apoptosis of ovarian cancer cells over normal fallopian epithelial cells [10]. Further studies with SpiD3 (3-carbon-linker SpiD) revealed that shorter linker lengths correlated with more potent inhibition of malignant cell growth [14]. Prompted by SpiD3’s impressive anti-leukemic activity in the NCI-60 cell line panel screen [14], we sought to assess SpiD3 in B-cell malignancies. Using preclinical models of CLL, we evaluated the anti-tumor properties of SpiD3 and investigated its molecular mechanism of action (MoA) by multi-omics and functional analyses. SpiD3 markedly inhibited malignant B-cell proliferation and suppressed NF-κB activation independent of TME-associated stimuli. Additionally, SpiD3 induced the UPR in CLL cells, resulting in prominent apoptosis and inhibition of protein synthesis. Concordantly, we witnessed a decreased tumor burden in SpiD3-treated mice with advanced leukemia. In ibrutinib-resistant CLL cells, SpiD3 sustained its anti-leukemic properties, further supporting the development of this compound for relapsed/refractory disease.

## MATERIALS AND METHODS

### Inhibitors and stimulants

SpiD3, SpiD7, and analog 19 were synthesized at University of Nebraska Medical Center (UNMC; Omaha, NE) following reported procedures [13–15]. Thapsigargin, cycloheximide, z-VAD(OMe)-FMK, ibrutinib, and JQ-1 were purchased from Cayman Chemical (Ann Arbor, MI). TPCA-1 was purchased from Sigma-Aldrich (Saint Louis, MO). All inhibitors were dissolved in DMSO. Stimulants used to mimic TME signals included 20 ng/mL TNF-α (Cayman Chemical), 500 ng/mL recombinant human (rh) sCD40 ligand (Peprotech; Cranbury, NJ), 50 ng/mL rhBAFF ligand (Peprotech), 10 μg/mL goat F(ab)^2^ anti-human IgM (Jackson ImmunoResearch; West Grove, PA), 3.2 μM CpG oligonucleotides 2006 (CpG; Integrated DNA Technologies; Coralville, IA), 1X lipopolysaccharide (LPS; Invitrogen; Carlsbad, CA) and 1X PMA/Ionomycin (BioLegend; San Diego, CA).

### Cell lines and primary CLL cell processing

Cell lines used in this study are HG-3, MEC1, MEC2, OCI-LY3, RI-1 (DSMZ; Braunschweig, Germany), DB, SUDHL6, Pfeiffer, RC, Jeko-1 (ATCC; Gaithersburg, MD), and 9-15c (RIKEN cell bank; Ibaraki, Japan). OSU-CLL cells [16] were provided by the Human Genetics Sample Bank of The Ohio State University (Columbus, OH). Ibrutinib-resistant HG-3 cells were generated by prolonged exposure to increasing concentrations of ibrutinib [17]. HG-3, RI-1, OSU-CLL, MEC1, MEC2, DB, SUDHL6, RC, Jeko-1, and 9-15c cell lines were cultured in RPMI-1640 with 2 mM L-glutamine (Sigma-Aldrich; St. Louis, MO), supplemented with 100 U/mL penicillin/100 μg/mL streptomycin (P/S, Sigma-Aldrich), and 10% heat-inactivated fetal bovine serum (hi-FBS, Avantor®; Radnor, PA). The Pfeiffer cell line was cultured in IMDM containing 25 mM HEPES and 2 mM L-glutamine, supplemented with P/S and 20% hi-FBS. The OCI-LY3 cell line was cultured in IMDM with 25 mM HEPES and 2 mM L-glutamine (Lonza; Switzerland), supplemented with P/S, 20% hi-FBS, and 1% β-mercaptoethanol (ThermoFisher Scientific; Waltham, MA). For all cell lines, a large stock was cryopreserved and used within 8 weeks after thawing. Prior to experimental use, cell line cultures were confirmed to be free of mycoplasma using the MycoAlert kit from Lonza.

CLL patient samples were obtained following informed consent under a protocol approved by the Institutional Review Board (IRB) of UNMC following the Declaration of Helsinki. CLL diagnosis was determined per iwCLL 2018 guidelines [18]. Patient characteristics are tabulated in Supplementary Table S1. Peripheral blood mononuclear cells (PMBCs) were isolated from patient blood using Lymphoprep™ density gradient centrifugation following manufacturer protocols (STEMCELL Tech; Vancouver, Canada) and confirmed to contain ≥ 90% CD19^+^/CD5^+^ cells before use. Patient-derived CLL samples were cultured in RPMI-1640 with 2 mM L-glutamine (Sigma-Aldrich) supplemented with P/S, and 10% hi-FBS. PBMCs from healthy donors were obtained from the Elutriation Core Facility (UNMC) per approved IRB protocol.

Primary murine tumor samples were isolated from moribund Eμ-Myc/TCL1 and Eμ-TCL1 spleens following institutional animal care guidelines at UNMC and confirmed to contain ≥ 90% CD19^+^/CD5^+^ cells before use. Eµ-Myc/TCL1 and Eμ-TCL1 spleen-derived malignant B-cells were cultured in RPMI-1640 + L-glutamine and supplemented with P/S, 10% hi-FBS, 0.055 mM β-mercaptoethanol, 10 mM HEPES (Sigma-Aldrich), 0.1 mM non-essential amino acid solution (Lonza), and 1 mM sodium pyruvate (Lonza).

### Cytotoxicity assays

Malignant B-cell lines (20-80,000 cells/well; 72 h), patient-derived CLL samples (0.7e^6^ cells/well; 48 h ± CpG), murine Eμ-Myc/TCL1 or Eμ-TCL1 lymphocytes (0.6e^6^ cells/well; 48 h ± PMA/Ionomycin) were treated with vehicle (DMSO) or increasing inhibitor concentrations (single or combined). Cell proliferation was evaluated using CellTiter 96® Aqueous MTS assay (Promega; Madison, WI) as previously described [17]. GraphPad Prism v9.4.1 (GraphPad Software, Inc.; San Diego, CA) was used to calculate the half-maximal inhibitory concentration (IC_50_). To assess synergy, combination indices (CI) were calculated using CompuSyn [19].

Cell viability and apoptosis were measured using Annexin V-FITC/propidium iodide (PI) assay kit (Leinco Technologies; Fenton, MO) per manufacturer’s protocol. For stromal co-cultures, 9-15c stromal cells (15,000 cells/well) were seeded in 48-well plates two days before adding patient-derived CLL cells (1e^7^ cells/mL) and subject to Annexin V/PI staining after 48 h inhibitor treatment.

### Immunoblot assays

Total cell protein was extracted using protein lysis buffer (20 mM Tris pH 7.4, 150 mM NaCl, 1% Igepal CA-630, 5 mM EDTA) containing protease and phosphatase inhibitor cocktails and phenylmethyl sulfonyl fluoride (Sigma-Aldrich). BCA protein analysis (ThermoFisher Scientific) was used to determine equal concentrations of protein for each sample lysate. Samples were then boiled for 5 min at 95°C in loading dye containing sodium dodecyl sulfate (SDS), separated by 1.5 mm SDS-PAGE gels, and then transferred onto nitrocellulose membranes using the Trans-Blot Turbo Transfer System (Bio-rad; Hercules, CA). Membranes were incubated overnight in primary antibody, washed, and incubated with HRP-conjugated anti-rabbit Ig or anti-mouse Ig (Cell Signaling Technology; Danvers, MA) for 1 h. Membranes were then visualized on the ChemiDoc Imaging System (Bio-rad) following development with WesternBright^TM^ ECL or Sirius (Advansta; San Jose, CA) according to the manufacturer’s instructions. Primary antibodies used are listed in Supplementary Table S2.

### RNA-sequencing and data analysis

RNA was extracted from OSU-CLL cells (1.5e^6^ cells/mL) treated with DMSO or SpiD3 (1, 2 μM; 4 h) using the miRNeasy Mini Kit (Qiagen; Hilden, Germany) per manufacturer instructions and processed using the Universal Plus mRNA-Seq with NuQuant kit (Tecan; Mannedorf, Switzerland). RNA quality was assessed on the Fragment Analyzer Automated CE System (Advanced Analytical Technologies, Inc; Ames, IA) and RNA (250 ng) was sequenced on Illumina NextSeq550 (San Diego, CA) at the UNMC Genomics Core. Sequenced reads were mapped using HISAT2 (v2.1.0) [20] to genome build hg38, guided by ENSEMBL GRCh38.99 transcript annotations. FPKM normalized values were obtained by Stringtie (v2.1.1) [21].

Differential gene expression was performed using limma [22]. Volcano analysis was performed using EnhancedVolcano (v1.2.0) [23]. Genes with *FDR*<0.05 and |Log_2_ FC|>0.5 were considered significant. The top 500 most significantly upregulated or downregulated genes were analyzed using the Molecular Signatures Database (MSigDB; v7.5.141-43) [24]. Scale-free weighted signed gene co-expression networks were constructed with the weighted gene co-expression network analysis (WGCNA) package [25] using the top 75% most variably expressed genes (n = 15005) according to their standard deviation. Detailed methodology is found in Supplementary Materials.

### Cell cycle analysis

OSU-CLL cells (1e^6^ cells/mL), synchronized overnight with aphidicolin (20 μg/mL, Sigma), were treated with inhibitors (48 h) and fixed with cold absolute ethanol (1 h). Fixed cells were stained with PI/Triton X-100/Rnase A solution as previously described [26] for flow cytometry analysis.

### Reactive oxygen species (ROS) detection

OSU-CLL cells (1e^6^ cells/mL), pretreated for 1 h with 5 mM N-acetylcysteine (Cayman Chemical), were incubated for 24 h with SpiD3 and stained using the ROS-detection cell-based assay kit (DCFDA; Cayman Chemical) per manufacturer’s protocol. Helix NIR (BioLegend) was used for live/dead discrimination prior to flow cytometry analysis. Fold change in median fluorescence intensity (MFI) was calculated as treatment group MFI / control MFI.

### Unfolded protein response (UPR) determination

Following treatment with SpiD3 or thapsigargin, CLL cell lines (1e^6^ cells/mL; 4 h) or CpG-stimulated patient-derived CLL samples (1e^7^ cells/mL; 24 h) were stained for 30 min with 50 μM thiol probe, tetraphenylethene maleimide (TPE-NMI), to assess cellular levels of SEC residues (unfolded/misfolded protein load) [27]. Helix NP™ NIR (BioLegend) was added for live/dead discrimination prior to flow cytometry analysis.

### Protein synthesis Click-iT assay

HG-3 cells (1e^6^ cells/mL) or CpG-stimulated patient-derived CLL samples (1e^7^ cells/mL) treated with inhibitors (24 h) or cycloheximide (30 min) were processed using the Protein Synthesis Assay Kit (Cayman Chemical) per manufacturer’s protocol. Fixable Zombie NIR (BioLegend) was used to gate live cells for flow cytometry analysis.

### Click chemistry

Following alkyne-tagged analog 19 treatment (10 μM; 2 h), protein lysates from OSU-CLL cells (1e^6^ cells/mL) were clicked to TAMRA biotin-azide and incubated with streptavidin agarose resin (Click Chemistry Tools; Scottsdale, AZ) to isolate biotin-alkyne-tagged proteins. For mass spectrometry analysis, the resin underwent trypsin digestion and peptides were run on an Orbitrap Fusion™ Lumos™ mass spectrometer (ThermoFisher Scientific) at the UNMC Proteomics Core and analyzed using Proteome Discoverer (ThermoFisher Scientific, v2.2). Pathway analysis was performed using EnrichR [28]. Biotin-alkyne-tagged proteins and their corresponding input lysates were subjected to immunoblotting for validation. Detailed methodology is found in Supplementary Materials.

### Murine Studies

All animal experiments were approved by the Institutional Animal Care and Use Committee (IACUC) at UNMC. Equal numbers of male and female Eμ-TCL1 transgenic mice [29] (median age = 10.2 mo) with evident leukemia in the blood (median leukemia burden = 51.8% CD45^+^/CD19^+^/CD5^+^ lymphocytes) were randomized to receive either SpiD3 prodrug (SpiD3_AP, 10 mg/kg) or vehicle equivalent (50% PEG400, 10% DMSO, and 40% water) daily via tail vein intravenous (IV) injection for 3 days (n = 6 mice/treatment arm). At study end (∼3 h after the last dose), mice were anesthetized with isoflurane (VetOne; Boise, ID) for tissue collection (blood and spleen) and downstream analyses. Whole blood was incubated (20 min at 4°C) with anti-CD45-APC, anti-CD3-PE/Cy7, anti-CD5-FITC (BioLegend) and anti-CD19-PE (BD Biosciences; Franklin Lakes, NJ) and then lysed using RBC Lysis Buffer (BioLegend) per manufacturer protocol before flow cytometry analysis. Harvested spleens were homogenized into a single cell suspension by passing through a 70 μm filter, subjected to RBC lysis, and used for downstream studies. Cells (∼2e^6^) were suspended in PBS/5% hi-FBS (100 uL) and incubated with anti-CD3-BUV737, anti-CD19-BUV395 (BD Horizon), anti-CD5-BV711, and anti-CD45-AF700 (BioLegend) for flow cytometric analysis. For immunoblot analysis, spleen-derived B-cells were selected using the EasySep™ Mouse Pan-B Cell Isolation Kit (STEMCELL Tech).

### Flow cytometry

Flow cytometry was performed on a LSRII (BD Biosciences), LSRFortessa X-50 (BD Biosciences), or NovoCyte 2060R (Agilent; Santa Clara, CA) cytometer and analyzed using NovoExpress v1.3.0 (Agilent) or Kaluza v2.1 (BD Biosciences).

### Statistics

Data are reported as mean ± standard error of the mean (SEM). Unpaired t-tests were used to compare two groups and one-way ANOVA with Dunnett’s multiple comparison test was used for comparing more than two groups using GraphPad Prism v9.4.1. *P* < 0.05 was considered significant.

## RESULTS

### SpiD3 displays cytotoxicity in B-cell malignancies

We first compared the anti-proliferative properties of analog 19 (monomer), SpiD3 (3-carbon-linker SpiD), and SpiD7 (7-carbon-linker SpiD) in CLL cell lines (Fig. 1A and B). Since SpiD3 was more potent than SpiD7 and analog 19 in HG-3 cells (∼10-fold) and OSU-CLL cells (∼5-fold; Fig. 1B), it was selected for further evaluation in this study. SpiD3 also exhibited anti-proliferative effects in chemotherapy resistant, *TP53* mutant MEC1 and MEC2 CLL cell lines [30] (IC_50_ = 0.5 μM; Fig. 1C). We next extended our evaluation of SpiD3 to a panel of lymphoma B-cell lines (Fig. 1C). Strikingly, SpiD3 also displayed high potency at sub-micromolar concentrations in aggressive mantle cell lymphoma and double-hit/triple-hit (DH/TH) lymphoma B-cells (IC_50_ < 0.9 μM). SpiD3 treatment induced apoptosis as indicated by increased Annexin V^+^ cells (Fig. 1D) and PARP cleavage (Fig. 1E) in CLL cell lines. SpiD3-mediated OSU-CLL cytotoxicity (Fig. 1F) and PARP cleavage (Fig. 1G) were partially reversed with the pan-caspase inhibitor, z-VAD-FMK.

**Figure 1:**
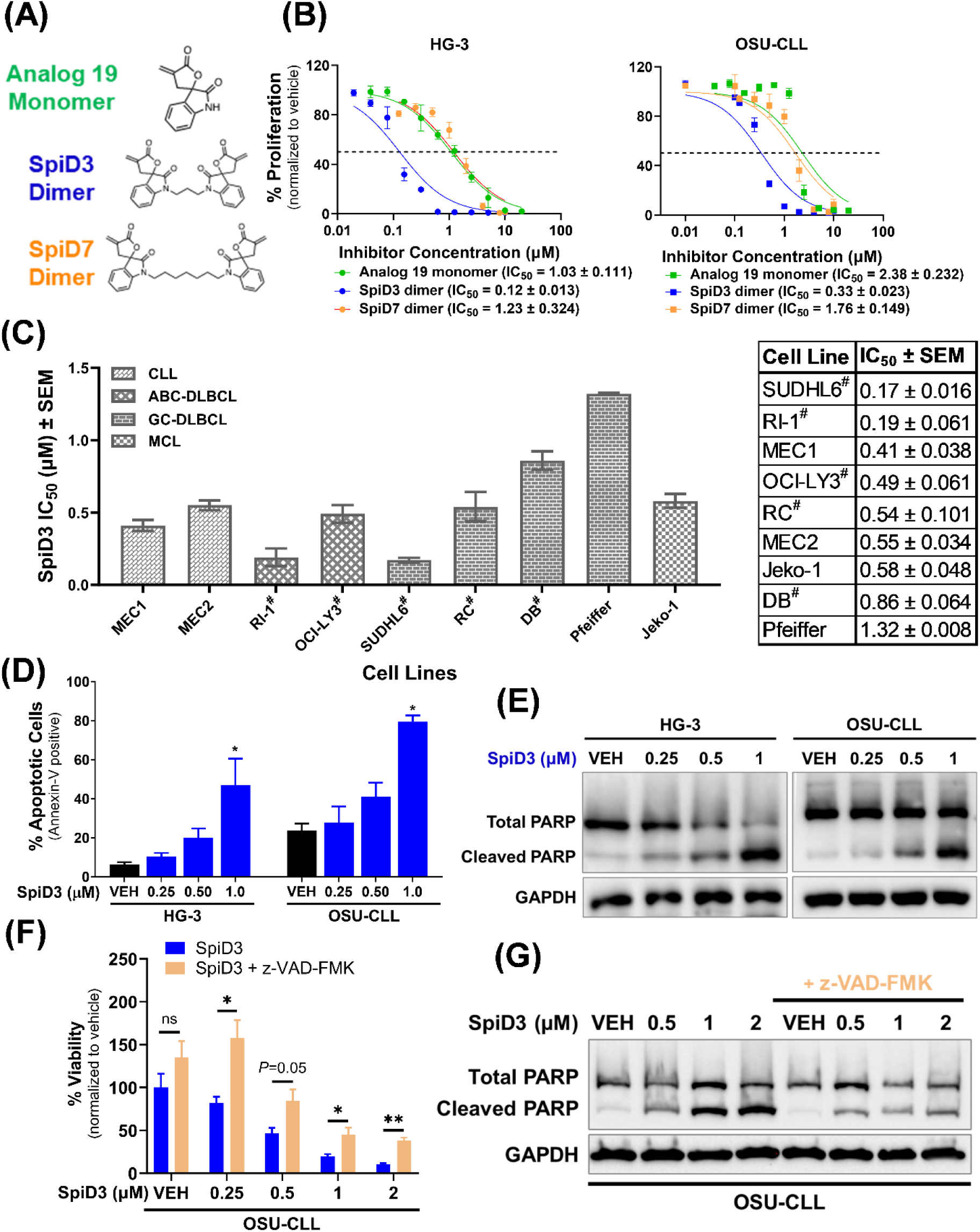
SpiD3 inhibits proliferation and induces apoptosis in malignant B-cells. **(A)** Chemical structures of analog 19 (monomer), SpiD3 and SpiD7 (dimers of analog 19). Full synthesis details reported elsewhere. **(B)** Proliferation of CLL cell lines, HG-3 and OSU-CLL, was determined by MTS assay following treatment with increasing concentrations of analog 19, SpiD3, and SpiD7 for 72 h (n = 3-4 independent experiments/cell line). **(C)** The IC_50_ values (mean ± SEM) of a panel of B-cell malignancy cell lines following SpiD3 treatment (72 h) was determined by MTS assay (n = 3-5 independent experiments/cell line). CLL: chronic lymphocytic leukemia, ABC-DLBCL: activated B-cell-like diffuse large B-cell lymphoma, GC-DLBCL: germinal-center-like diffuse large B-cell lymphoma, MCL: mantle cell lymphoma. ^#^Indicates a double-hit lymphoma or a triple-hit lymphoma. On the right is a table of the IC_50_ values (mean ± SEM). **(D)** Percent apoptosis of HG-3 and OSU-CLL cell lines was determined by Annexin V/PI viability assay following 24 h treatment with SpiD3 (n = 4 independent experiments/cell line). **(E)** Representative immunoblot analyses of total and cleaved PARP in OSU-CLL and HG-3 cells following 4 h treatment with SpiD3 or DMSO vehicle (VEH). GAPDH serves as the loading control (n = 6 independent experiments/cell line). **(F)** MTS assay of OSU-CLL cells (n = 3 independent experiments) treated for 24 h with increasing concentrations of SpiD3 (blue) with or without co-current z-VAD-FMK treatment (50 μM, tan). **(G)** Representative immunoblot analyses of total and cleaved PARP in OSU-CLL cells treated with SpiD3 for 4 h in the presence or absence of z-VAD-FMK pretreatment (1 h, 50 μM). GAPDH served as the loading control (n = 4 independent experiments). Error bars and IC_50_ values are shown as mean ± SEM. Asterisks denote significance *vs*. VEH: \**P* < 0.05, \*\**P* < 0.01.

### SpiD3 induces a unique transcriptional and translational program in CLL cells

Using a comprehensive multi-omics approach, we explored the MoA of SpiD3 in CLL. SpiD3-treated OSU-CLL cells were subjected to RNA-sequencing (Fig. 2; Supplementary Fig. S1) and tandem mass tag (TMT) proteomics (Supplementary Fig. S2) analyses. Transcriptional analysis revealed SpiD3 modulated over 6500 genes (*FDR* < 0.05) following a 4 h exposure to 2 μM SpiD3 (Fig. 2A). On the protein level, 132 proteins (*P* < 0.05) were modulated following a 24 h exposure to 1 μM SpiD3 (Supplementary Fig. S2A and S2B).

**Figure 2:**
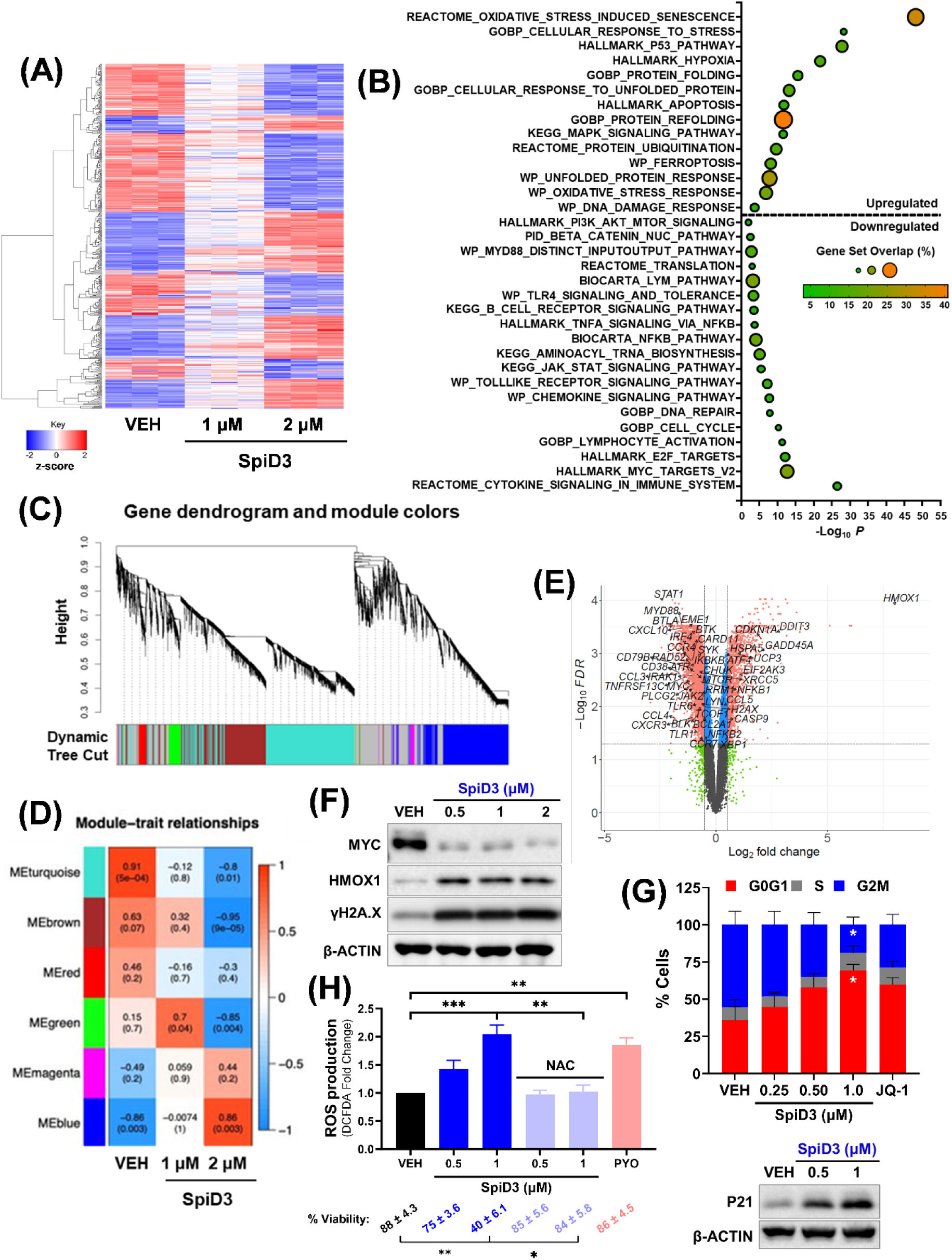
SpiD3 modifies transcriptional profiles and subverts oncogenic pathways in CLL cells. **(A-E):** RNA-sequencing of OSU-CLL cells treated with SpiD3 (1, 2 μM; 4 h) or equivalent DMSO vehicle (VEH; n = 3 independent experiments). (A) Hierarchical clustering of the top 500 differentially expressed genes (DEGs) in SpiD3-treated cells (*FDR* < 0.05). Red indicates increased gene expression (0 < z-score < 2), and blue indicates decreased expression (-2 < z-score < 0). **(B)** Gene set enrichment analysis of the top 500 DEGs in 2 μM SpiD3-treated cells. **(C)** Gene dendrogram and colors of WGCNA-identified modules for the top 500 DEGs (*P* = 0.05) in SpiD3-treated cells (1, 2 μM). **(D)** Heatmap of correlation between WGCNA module and the indicated treatment conditions. Each heatmap cell displays the correlation coefficient (top) and corresponding *P*-value (bottom). **(E)** Volcano plot of SpiD3-treated cells (2 μM) with select CLL-relevant genes labeled. Genes meeting both the statistical significance (*FDR* < 0.05) and fold-change (|Log_2_ FC|> 0.5) were used for downstream analysis (red). Genes meeting only statistical significance (blue), only fold-change (green), or neither threshold (grey) are shown for comparison. **(F)** Representative immunoblots (n = 3 independent experiments) of the indicated proteins in SpiD3-treated OSU-CLL cells (4 h). **(G)** Cell cycle analysis of OSU-CLL cells treated with increasing amounts of SpiD3 (48 h). BET inhibitor, JQ-1 (1 μM), served as a positive control for cell cycle arrest (n = 5 independent experiments). Insert depicts a representative immunoblot for P21 expression following SpiD3 treatment (24 h; n = 3 independent experiments). β-ACTIN served as the loading control. **(H)** OSU-CLL cells were pretreated with 5 mM N-acetylcysteine (NAC, 1 h) followed by SpiD3 or VEH (24 h). Pyocyanin (PYO, 1 mM) served as a control ROS inducer (n = 3 independent experiments). Percent viability per condition is denoted below. Data are represented as mean ± SEM. Asterisks denote significance *vs.* VEH: \**P* < 0.05, \*\**P* < 0.01, \*\*\**P* < 0.001.

Gene set enrichment analysis (GSEA) of our multi-omics data revealed SpiD3 influenced various pathways highly relevant to CLL survival, proliferation and TME interplay such as “BCR signaling”, “NF-κB signaling”, “cytokine signaling”, “E2F and MYC targets”, “oxidative stress”, “DNA damage”, “P53 pathway” and modes of cell death like “apoptosis” (Fig. 2B; Supplementary Fig. S1B; Supplementary Fig. S2C). In addition, the pathway analyses highlighted increased UPR and reduced protein translation in SpiD3-treated CLL cells (Fig. 2B; Supplementary Fig. S1B; Supplementary Fig. S2C). SpiD3 mediated transcriptional changes to *HMOX1*, *γH2AX*, and *MYC*, consistent with corresponding changes in protein expression (Fig. 2E and F; Supplementary Fig. S1A; Supplementary Fig. S2A and S2B).

To identify modules of highly correlated genes and assess their relationships and biological functions in SpiD3-treated CLL cells, we performed weighted gene co-expression network analysis (WGCNA). WGCNA identified 28 modules, 6 reflected the most characteristic changes following 2 μM SpiD3 treatment (Fig. 2C and D). Other modules containing less significantly altered genes are represented in grey. Turquoise, brown, green, and red colored modules negatively correlated with SpiD3 treatment. Genes in these modules were related to “DNA repair”, “tRNA modification”, “PI3K signaling”, “ER protein trafficking”, “chemokine signaling”, “lymphocyte activation”, and “cell cycle”, suggesting SpiD3 negatively impacts CLL protein processing, influences CLL-TME crosstalk, and induces cell cycle arrest. Conversely, the blue and magenta modules were positively correlated with SpiD3 treatment and consisted of genes related to “stress-activated kinase signaling”, “DNA damage”, and “ER-associated degradation”, suggesting SpiD3 heightens ER stress, which can trigger UPR induction and subsequently attenuate CLL cell growth and survival.

To support our multi-omics findings, we next evaluated SpiD3-mediated effects on CLL cell cycle progression, ROS production, and chemotaxis. SpiD3-treated CLL cells accumulated in the G0/G1 phase with a corresponding reduction in the G2/M population (Fig. 2G) comparable to JQ-1, a potent inducer of cell cycle arrest [26]. This coincided with upregulation of P21 expression in SpiD3-treated OSU-CLL cells (Fig. 2G). SpiD3 also induced ROS production in CLL cells comparable to pyocyanin (control ROS inducer). Intriguingly, pretreatment with the antioxidant, N-acetylcysteine, not only mitigated SpiD3-induced ROS production, but also attenuated SpiD3-mediated cytotoxicity (Fig. 2H). Comparable to ibrutinib, SpiD3 reduced OSU-CLL cell migration toward CXCL-12, a chemokine secreted by bone marrow stromal cells [31] (Supplementary Fig. S3A).

### SpiD3 induces the UPR and inhibits protein translation in CLL

Given our recent study demonstrating SpiD7 induces the UPR through crosslinking cellular proteins in ovarian cancer cell lines [10], we sought to confirm if SpiD3 generates unfolded/misfolded proteins in CLL cells. We used TPE-NMI; a thiol probe that fluoresces once bound to SEC groups on unfolded proteins [27]. SpiD3 dose-dependently increased TPE-NMI fluorescent intensity (Fig. 3A), implying an amplified burden of unfolded proteins in SpiD3-treated CLL cells. Accordingly, SpiD3 induced XBP1 splicing (Fig. 3B, blue arrow), comparable to thapsigargin (UPR-agonist), indicating IRE1α activation. We additionally observed a PERK band shift indicative of its phosphorylation (Fig. 3B, green arrow), accompanied by marked induction of eIF2α phosphorylation (Fig. 3B) in SpiD3-treated CLL cell lines. *DDIT3* (*CHOP*) and *ATF4* contain short open reading frames in their 5’ untranslated regions which allows them to be preferentially translated when eIF2α is phosphorylated [8]. As expected, SpiD3 treatment increased mRNA and protein expression of ATF4 and CHOP (Fig. 3B; Supplementary Fig. S3B).

**Figure 3:**
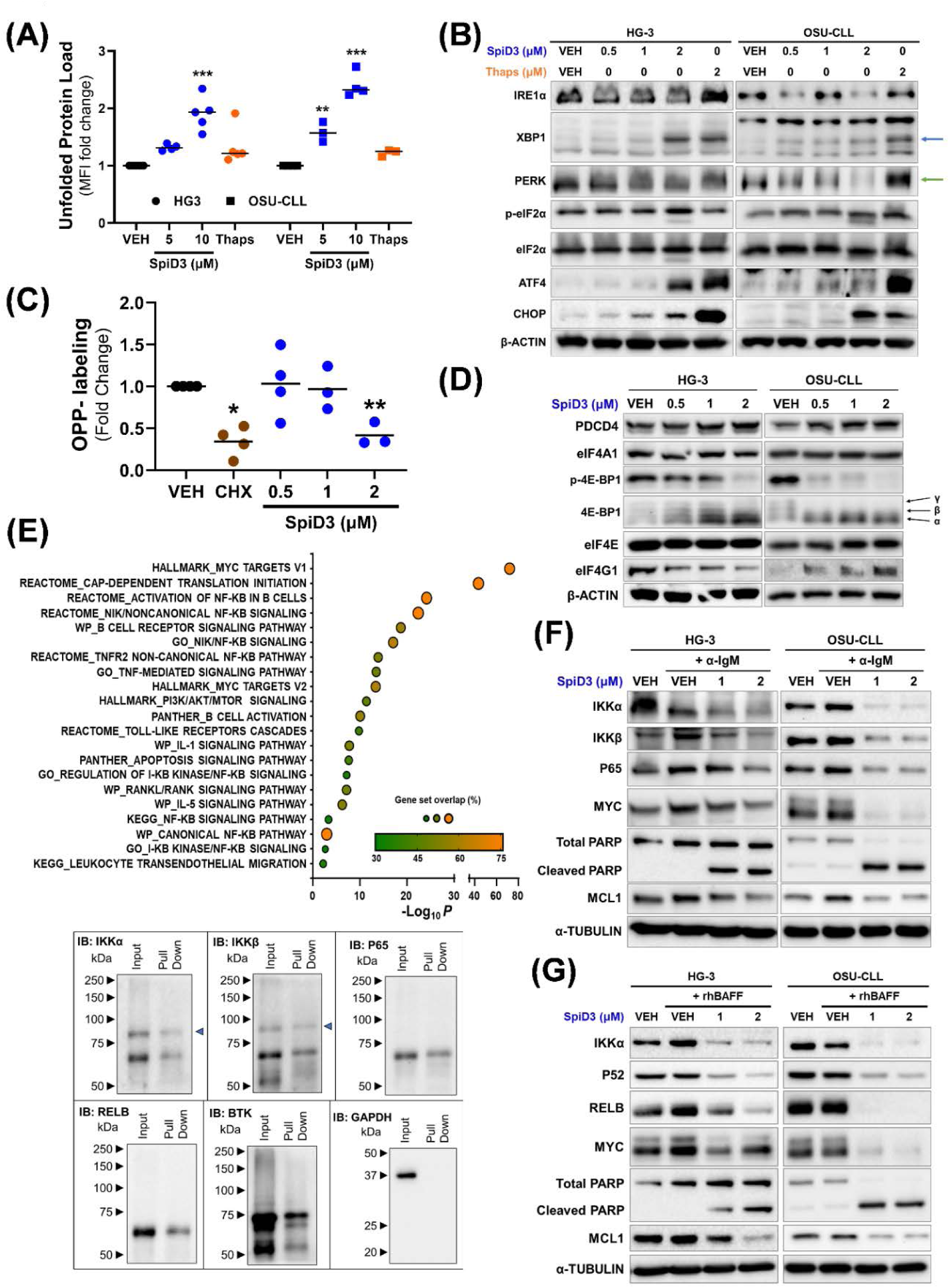
SpiD3 induces the unfolded protein response and modulates CLL survival factors independent of TME stimuli. **(A)** HG-3 and OSU-CLL cells were treated for 4 h with SpiD3 (5, 10 μM), thapsigargin (Thaps; 10 μM), or equivalent DMSO vehicle (VEH) and then incubated with TPE-NMI dye to probe for unfolded proteins. Data are represented as fold change in TPE-NMI median fluorescent intensity (MFI) compared to VEH (n = 3-5 independent experiments/cell line). **(B)** Representative immunoblot analyses of IRE1α, XBP1, PERK, ATF4, CHOP, p-eIF2α (Ser51), and total eIF2α in HG-3 and OSU-CLL cells treated with VEH, SpiD3 (0.5-2 μM), or Thaps (2 μM) for 4 h (n = 4-5 independent experiments/cell line). β-ACTIN served as the loading control. Blue arrow: spliced XBP1, green arrow: PERK shift. **(C)** HG-3 cells were treated with VEH (24 h), SpiD3 (0.5-2 μM; 24 h), or cycloheximide (CHX; 50 μg/mL; 30 min) and then incubated with O-propargyl-puromycin (OPP) for 30 min. Data are represented as fold change in OPP MFI compared to VEH (n = 3-4 independent experiments). **(D)** Representative immunoblot analyses of PDCD4, eIF4A1, p-4E-BP1 (Ser65), total 4E-BP1, eIF4E, and eIF4G1 levels in HG-3 and OSU-CLL cells treated with VEH or SpiD3 (0.5-2 μM) for 4 h (n = 4 independent experiments/cell line). β-ACTIN served as the loading control. Black arrows indicate the three isoforms of 4E-BP1. **(E)** OSU-CLL cells were incubated with the alkyne-tagged analog 19 (10 μM) for 2 h. Cell lysates were clicked with TAMRA-biotin and biotin-tagged-19-bound-proteins were isolated using streptavidin agarose beads, trypsinized, and then evaluated via mass spectrometry. **Top:** Pathway enrichment (EnrichR) analysis of similar proteins found in at least 2 of the 3 biological replicates. **Bottom:** Representative immunoblot analysis of biotin-alkyne-tagged proteins, with their corresponding input lysates for IKKα, IKKβ, P65, RELB, BTK, and GAPDH. The blue arrow indicates IKKα and IKKβ. **(F-G):** Immunoblot analysis of the indicated proteins in whole cell lysates of HG-3 and OSU-CLL cells treated with SpiD3 (1, 2 μM) for 4 hr and co-currently stimulated with **(F)** α-IgM (10 μg/mL) or **(G)** rhBAFF ligand (50 ng/mL). α-TUBULIN served as the loading control (n = 4 independent experiments/cell line). Asterisks denote significance vs. VEH: \**P* < 0.05, \*\**P* < 0.01, \*\*\**P* < 0.001.

PERK-mediated phosphorylation of eIF2α inhibits global protein translation by blocking ternary complex (i.e., eIF2-GTP, tRNA-Met) formation, a critical rate-limiting step in mRNA translation [32]. Consistent with the suppressed protein synthesis following UPR induction observed above, our multi-omics analyses revealed SpiD3 impacts protein translation pathways (e.g., decreased *mTOR, RRM1*, and RSP9 expression; Fig. 2E; Supplementary Fig. S2B). To further evaluate if SpiD3 inhibits global protein synthesis in leukemic cells, we incubated SpiD3-treated HG-3 cells with O-propargyl-puromycin (OPP), which incorporates into nascent polypeptide chains. SpiD3 (2 μM) significantly reduced OPP signal akin to cycloheximide, a control protein translation inhibitor (Fig. 3C).

Another rate-limiting step in protein translation initiation is eIF4F complex (eIF4A1, eIF4E, eIF4G1) assembly [32]. The expression of eIF4G1 was moderately reduced with SpiD3 only in HG-3 cells but, surprisingly, neither eIF4E nor eIF4A1 expression was altered by SpiD3 (Fig. 3D). However, in both HG-3 and OSU-CLL cells, SpiD3 modified the expression of key regulators of protein translation initiation, namely tumor suppressor programmed cell death 4 (PDCD4) and eukaryotic initiation factor 4E-binding protein 1 (4E-BP1), known protein inhibitors of eIF4A1 and eIF4E, respectively [32]. SpiD3 spared PDCD4 degradation, suggesting suppressed protein translation through depletion of available eIF4A1 (Fig. 3D). Furthermore, studies have shown 4E-BP1 competes with eIF4G1 in binding to eIF4E, and the hypo-phosphorylated (α) state of 4E-BP1 has a higher binding affinity to eIF4E than eIF4G1, thereby blocking eIF4E from the eIF4F complex [33]. Alternatively, 4E-BP1 hyper-phosphorylation facilitates its release from eIF4E, allowing cap-dependent translation to proceed [34]. SpiD3 inhibited 4E-BP1 phosphorylation at Ser56, thereby causing its accumulation in the hypo-phosphorylated (α) state (Fig. 3D). To determine if SpiD3 disrupts the eIF4G1:eIF4E interaction, we utilized agarose-immobilized m^7^GTP cap analogs [33] to capture eIF4E and its binding partners (eIF4G1, 4E-BP1). Following SpiD3 treatment, 4E-BP1 displaced eIF4G1 from eIF4E in the cap-bound fraction, comparable to that observed in serum-starved CLL cells (Supplementary Fig. S4). Overall, these data suggest SpiD3 impairs cap-dependent protein translation through inhibiting ternary complex and eIF4F complex assembly.

### SpiD3 targets disease-relevant proteins in CLL independent of TME stimuli

As previously reported, SpiD3 and analog 19 covalently bind to SEC residues through the Michael acceptor moiety [14, 15]. Using alkyne-tagged analog 19 [15], we adopted a click-chemistry mass spectrometry approach to identify proteome-wide targets of SpiD3 in CLL. Pathway enrichment analysis revealed that analog 19 targeted pathways related to protein translation, NF-κB, and BCR signaling in OSU-CLL cells (Fig. 3E). Immunoblot analyses validated interactions of analog 19 with key BCR (BTK) and NF-κB (P65, IKKβ, IKKα, RELB) pathway proteins (Fig. 3E), indicating SpiD3 affects key CLL survival pathways and disrupts both canonical and non-canonical NF-κB signaling. Activation of the NF-κB pathway in CLL promotes transcription of genes regulating inflammatory, proliferative, and anti-apoptotic processes through nuclear translocation [2]. SpiD3 treatment of TNF-α stimulated CLL cells resulted in marked inhibition of P65 (canonical) and RELB (non-canonical) nuclear translocation, whereas TPCA-1 (control NF-κB inhibitor) only blocked P65 nuclear translocation (Supplementary Fig. S5A).

Since TME interactions fuel various survival axes and downstream NF-κB activation in CLL cells [3], we sought to evaluate the anti-leukemic effects of SpiD3 under stimulation by TME mimetics. Under anti-IgM stimulation (BCR activation), SpiD3 decreased expression of canonical NF-κB pathway proteins, IKKα, IKKβ, P65 (Fig. 3F). SpiD3 retained its inhibitory effects on non-canonical pathway proteins (IKKα, RELB) under B-cell-activating factor (BAFF) receptor stimulation, which mimics interactions with nurse-like [6] or bone-marrow stromal [31] cells (Fig. 3G). Remarkably, SpiD3 attenuated NF-κB activation in the presence of soluble CD40 ligand (sCD40L), a known activator of both canonical and non-canonical NF-κB pathways [35] (Supplementary Fig. S5B), suggesting SpiD3 disrupts supportive CD40L-dependent T-cell mediated interactions in CLL [35]. In the presence of each TME mimetic, SpiD3 induced PARP cleavage, decreased oncogenic MYC expression, and reduced levels of MCL1, an anti-apoptotic protein contributing to CLL drug resistance [7] (Fig. 3F and G; Supplementary Fig. S5B).

### SpiD3 displays potent anti-tumor effects in patient-derived CLL samples

Prompted by the impressive anti-tumor properties of SpiD3 in malignant B-cell lines, we set out to validate these properties in patient-derived CLL samples including those harboring features of poor outcomes and/or high-risk disease (e.g., unmutated *IGHV,* deletion 17p, deletion 11q, complex karyotype) [36]. Patient characteristics are tabulated in Supplementary Table S1. Because other NF-κB inhibitors have failed to induce anti-tumor cytotoxic effects within protective tumor niches [37], we first evaluated the efficacy of SpiD3 in the presence of stromal cell support. SpiD3 dose-dependently reduced viability of patient-derived CLL cells co-cultured with bone marrow stromal cells (Fig. 4A). Importantly, SpiD3 was non-toxic to the bystander stromal cells, indicating SpiD3-induced CLL cytotoxicity was not attributed to reduced stroma viability (Supplementary Fig. S6A). To mimic proliferative signals witnessed in CLL pseudofollicles, patient-derived CLL samples were stimulated ex vivo with CpG, a powerful TLR9 agonist [38]. Excitingly, SpiD3 impaired CpG-induced proliferation of primary CLL samples in a dose dependent manner (Fig. 4B) while sparing healthy donor lymphocytes (Supplementary Fig. S6B). In comparison, TPCA-1 (control NF-κB inhibitor) was cytotoxic toward both the healthy donor and CLL samples (Supplementary Fig. S6B).

**Figure 4:**
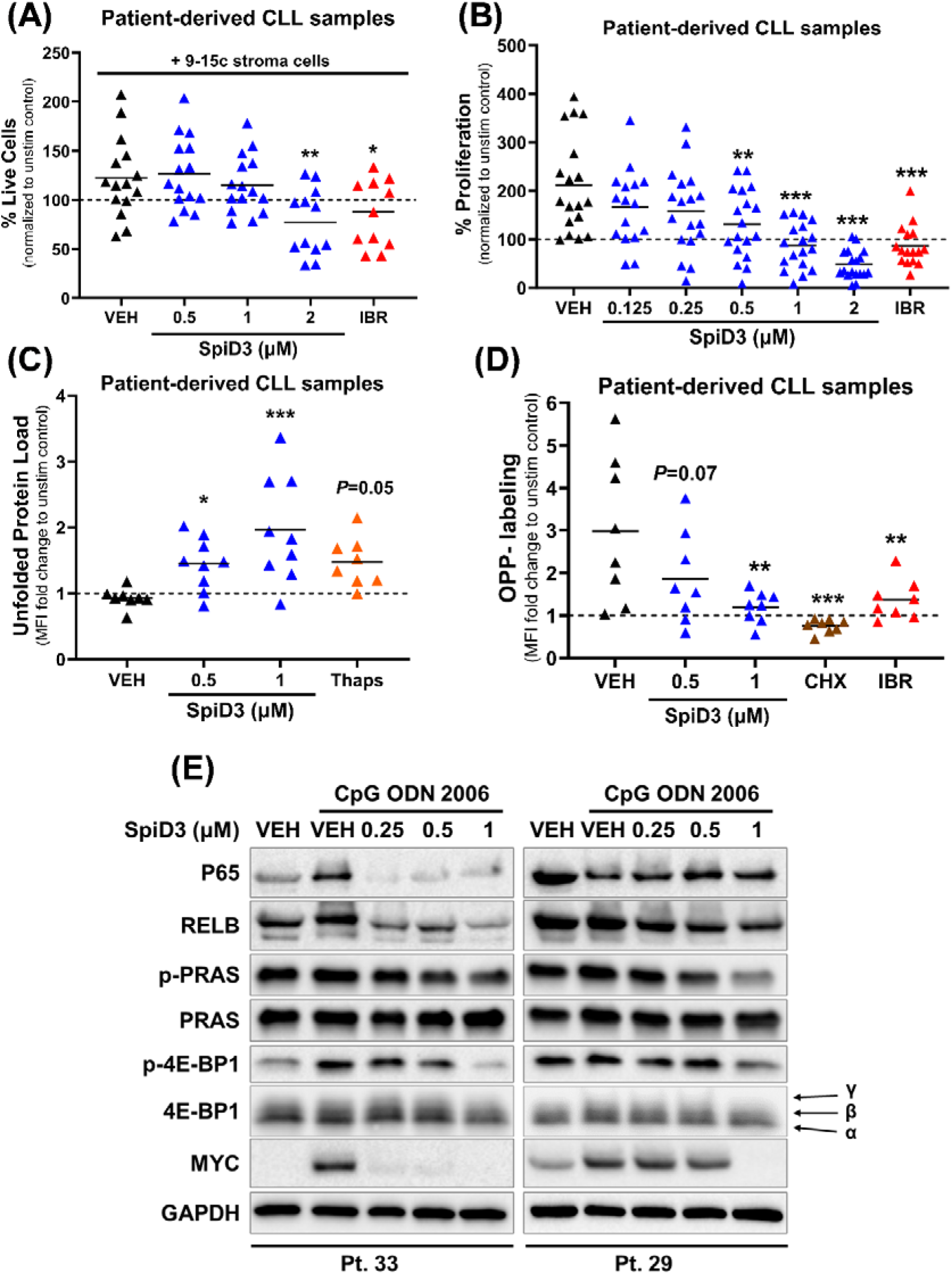
Anti-tumor effects of SpiD3 in patient-derived CLL samples. **(A)** Viability of patient-derived CLL samples (n = 11-15) co-cultured ex vivo with 9-15c mouse stroma cells for 48 h was determined by Annexin-V/PI staining. Ibrutinib (IBR; 1 μM) serves as a control anti-leukemic drug. Results are normalized to the unstimulated control (dashed line). **(B)** Following ex vivo treatment with SpiD3 (0.125-2 μM), the relative proliferation of patient-derived CLL samples (n = 16-18) under co-current CpG ODN 2006 stimulation (3.2 μM) was assessed via MTS assay (48 h). IBR (1 μM) was used as control anti-leukemic drug. Results are normalized to the unstimulated control (dashed line). **(C)** UPR induction in patient-derived CLL samples (n = 8) was evaluated following 24 h ex vivo treatment with SpiD3 or thapsigargin (Thaps; 1 μM) under co-current CpG stimulation (3.2 μM) via incubation with TPE-NMI dye. Data are represented as fold change in TPE-NMI median fluorescence intensity (MFI) compared to the unstimulated control (dashed line). **(D)** Protein synthesis in patient-derived CLL samples (n = 8) was assessed via OPP incorporation following a 24 h ex vivo treatment with SpiD3, cycloheximide (CHX; 50 μg/mL), or IBR (1 μM) under co-current CpG stimulation (3.2 μM). Data are represented as fold change in OPP-Alexa Fluor MFI compared to the unstimulated control (dashed line). **(E)** Representative immunoblot analyses of P65, RELB, p-PRAS (Thr246), total PRAS, p-4E-BP1 (Ser65), total 4E-BP1, and MYC protein in patient-derived CLL samples (n = 6) following a 24 h treatment with SpiD3 in the presence of CpG (3.2 μM). Black arrows indicate the three isoforms of 4E-BP1. GAPDH served as the loading control. Patient characteristics are tabulated in Supplementary file 1: Table S1. Asterisks denote significance *vs*. stimulated VEH: \**P* < 0.05, \*\**P* < 0.01, \*\*\**P* < 0.001.

To verify the MoA of SpiD3 in patient-derived CLL samples, we evaluated UPR induction and protein translation through flow cytometry based-assays. Remarkably, SpiD3 induced UPR in primary CLL cells (Fig. 4C), but not in the healthy donor B-cells (Supplementary Fig. S6C), suggesting that the higher basal UPR in CLL cells renders them more sensitive to agents like SpiD3 that can covalently modify SECs. Notably, SpiD3 treatment inhibited protein translation in primary CLL cells (Fig. 4D). Immunoblot analysis of SpiD3-treated primary CLL cells revealed reduced expression of NF-κB proteins (P65, RELB), cell survival signaling (p-PRAS/PRAS), and proliferation factors (MYC). This was accompanied by reduced phosphorylation of 4E-BP1 at Ser56 (Fig. 4E). These results indicate SpiD3 exerts a favorable cytotoxicity profile in primary CLL cells and inhibits protein synthesis via UPR overactivation, thereby presenting a unique means by which this novel agent elicits its anti-tumor properties.

### SpiD3 sensitizes CLL cells to ibrutinib and is effective in ibrutinib-resistant cells

BTK inhibitors like ibrutinib are efficacious in CLL, however, a growing number of patients progress on and/or develop resistance to ibrutinib [39]. To discern if SpiD3 can sensitize CLL cells to ibrutinib, we conducted combination studies to determine synergism between ibrutinib and SpiD3. In the aggressive unmutated *IGHV* HG-3 cell line [40], synergy was observed with 0.125 μM SpiD3 and 0.125 μM ibrutinib (Fig. 5A and B). Furthermore, strong synergy was displayed between 0.5 μM SpiD3 and 0.5 μM ibrutinib in patient-derived CLL cells (Fig. 5C and D).

**Figure 5:**
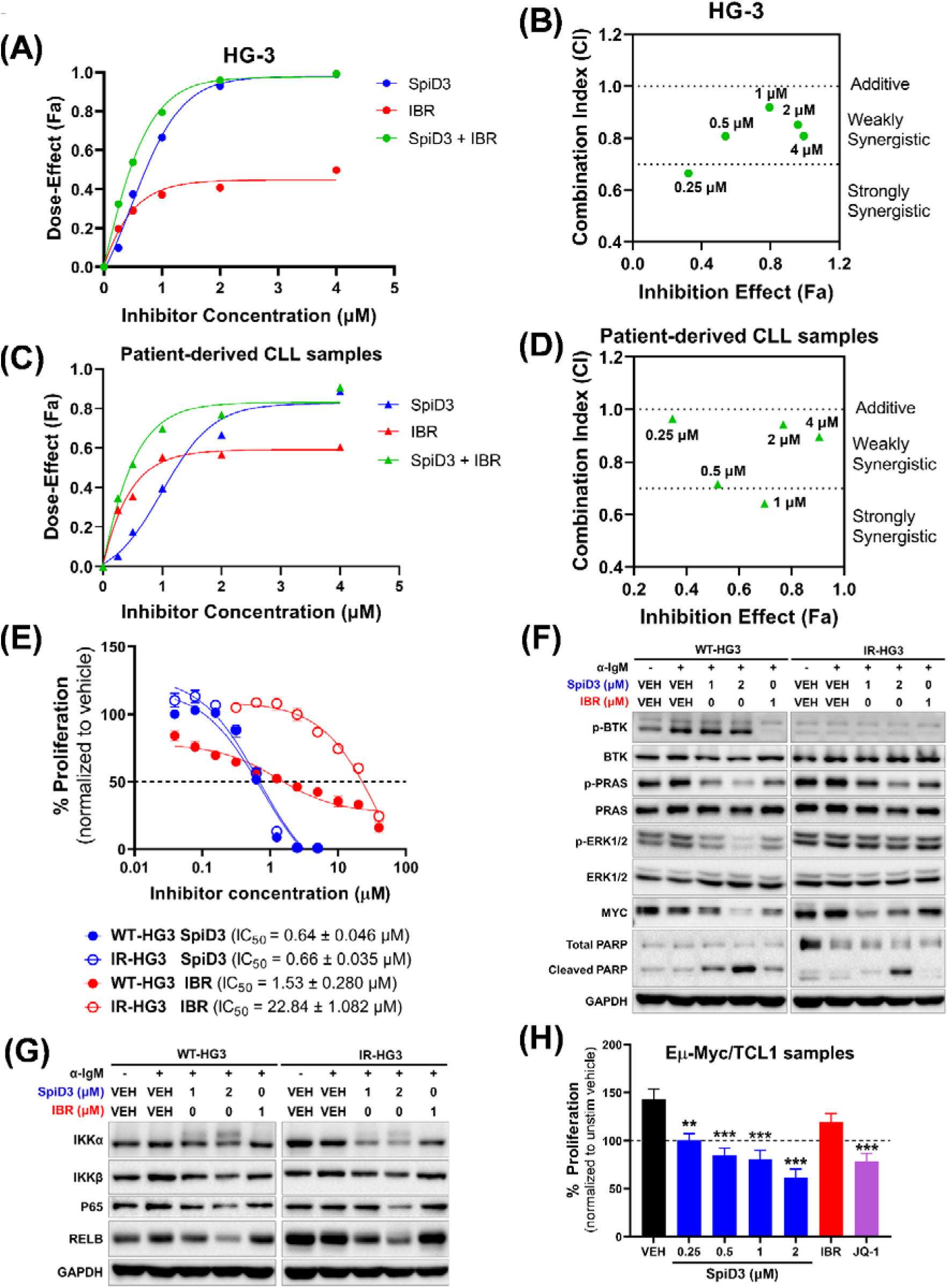
SpiD3 synergizes with ibrutinib and elicits cytotoxic effects in ibrutinib-resistant CLL cells. **(A-D):** Combination assays to test synergy between SpiD3 and ibrutinib (IBR; BTK inhibitor) in preclinical CLL models. HG-3 cells (**A-B**; n = 3 independent experiments) or CpG ODN 2006 (CpG; 3.2 μM)-stimulated patient-derived CLL cells (**C-D**; n = 6) were treated with SpiD3, IBR or both (1:1 ratio) for 72 h. MTS assay was performed to detect the dose-effect. **(A, C):** Dose-effect of the single drugs and their combination (1:1 ratio). **(B, D):** Combination index (CI) of the combined doses (0.5 μM SpiD3 + 0.5 µM IBR = 1 µM on graph). CI values were calculated using the Chou-Talalay method by the software Compusyn. CI values > 1 are antagonistic, CI values = 1 are additive, and CI values < 1 are synergistic. **(E)** Parental wild-type (WT) and ibrutinib-resistant (IR) HG-3 cell proliferation was assessed by MTS assay following treatment with increasing concentrations of SpiD3 or IBR (72 h; n = 6 independent experiments/cell line). IC_50_ values (mean ± SEM) are noted for each cell line. **(F-G):** Representative immunoblot analyses of p-BTK (Tyr223), total BTK, p-PRAS (Thr246), total PRAS, p-ERK1/2 (Thr202/Tyr204), total ERK1/2, MYC, PARP (total and cleaved), IKKα, IKKβ, P65, and RELB in WT- and IR-HG3 cells treated with SpiD3 (1-2 μM) or IBR (1 μM) for 4 h. BCR activation was induced by adding soluble α-IgM (10 μg/mL) for the last 15 min of treatment (n = 5 independent experiments). GAPDH served as the loading control. **(H)** Spleen-derived malignant B-cells from terminally diseased Eμ-Myc/TCL1 mice (n = 8) were stimulated ex vivo with 1X PMA/Ionomycin and treated with SpiD3 (0.25-2 μM), IBR (1 μM), or JQ-1 (1 μM) for 48 h. Proliferation was assessed via MTS assay and normalized to the unstimulated vehicle (dashed line). Asterisks indicate significance *vs.* stimulated vehicle. \**P* < 0.05, \*\**P* < 0.01, \*\*\**P* < 0.001.

Enhanced NF-κB activity has been reported in various ibrutinib-resistant B-cell malignancies [39]. Since SpiD3 sensitized CLL cells to ibrutinib, we sought to assess the effects of SpiD3 in ibrutinib-resistant HG-3 (IR-HG3) cells. Consistent with reported studies [17], the anti-proliferative effects of ibrutinib were attenuated in IR-HG3 cells. Remarkably, SpiD3 exhibited robust anti-proliferative effects in IR-HG3 cells with comparable IC_50_ to parental wild-type HG-3 (WT-HG3) cells (Fig. 5E). SpiD3 correspondingly decreased MYC expression, reduced PRAS and ERK1/2 phosphorylation, and induced PARP cleavage in both WT- and IR-HG3 cells. However, in IR-HG3 cells, ibrutinib was less effective in inhibiting PRAS and ERK1/2 activation, reducing MYC protein expression, or inducing PARP cleavage (Fig. 5F). In contrast to ibrutinib, SpiD3 also decreased IKKα, IKKβ, P65 and RELB expression in both cell lines, indicating its ability to target alternative survival mechanisms in ibrutinib-resistant disease (Fig. 5G). Finally, we evaluated SpiD3 ex vivo using primary tumor lymphocytes isolated from Eμ-Myc/TCL1 mice; an aggressive model of concurrent CLL and lymphoma [41]. Previous studies demonstrated that Eμ-Myc/TCL1 tumors are inherently resistant to ibrutinib treatment but responsive to novel agents like BET inhibitors [17, 26]. Remarkably, SpiD3 dose-dependently reduced Eμ-Myc/TCL1 lymphocyte proliferation comparable to JQ-1 (BET inhibitor), whereas ibrutinib was ineffective (Fig. 5H). These studies introduce SpiD3 as a promising agent to combat ibrutinib resistance, making it a viable pre-therapeutic lead for relapsed/refractory B-cell malignancies.

### SpiD3 reduces leukemia burden in Eμ-TCL1 mice

To establish the translational potential of spirocyclic dimers in CLL, we treated leukemic Eμ-TCL1 mice [29] with a prodrug of SpiD3 (SpiD3_AP). Reactive functional groups such as α, β-unsaturated systems found in the α-methylene-γ-butyrolactone of SpiD3 can be masked using prodrug strategies. Using our previously reported method [42], we have generated a dimethylamino prodrug of SpiD3 to improve stability in biological matrices for in vivo preclinical anti-tumor testing (Fig. 6A). We outline the detailed synthesis methodology of SpiD3_AP in the Supplementary Materials. Notably, SpiD3_AP has comparable anti-proliferative effects to SpiD3 in ex vivo treated Eμ-TCL1 spleen-derived lymphocytes (Fig. 6B) and the HG-3 cell line (Supplementary Fig. S7A). We initially evaluated microsomal stability to assess oral delivery as a potential route of administration. Although the prodrug, SpiD3_AP, exhibited an improved half-life compared to SpiD3 (Fig. 6C), the short half-life within liver microsomes promoted us to consider intravenous (IV) injection. The pharmacokinetic parameters of 10 mg/kg SpiD3_AP (Fig. 6D) further established IV administration and a collection time of 3 h after the final dose as the optimal parameters for this proof-of-concept study. Excitingly, a three-day SpiD3_AP treatment in Eµ-TCL1 mice resulted in a significant decrease in leukemia burden in the blood (Fig. 6E) and spleen (Fig. 6F) compartments when compared to vehicle-treated mice. Mouse body weight and bystander T-cells percentages were stable, indicating SpiD3_AP treatment was well-tolerated (Supplementary Fig. S7B and S7D). Immunoblot analysis of spleen-derived B-cells revealed a marked decrease in oncogenic MYC protein expression in SpiD3_AP-treated mice compared to vehicle-treated mice (Fig. 6G) further confirming the anti-proliferative potential of this novel therapeutic approach. This initial proof-of-concept study in Eµ-TCL1 mice with advanced disease demonstrates for the first time the efficacy of SpiD3_AP in CLL and advocates for continued in vivo exploration of SpiD prodrug candidates.

**Figure 6:**
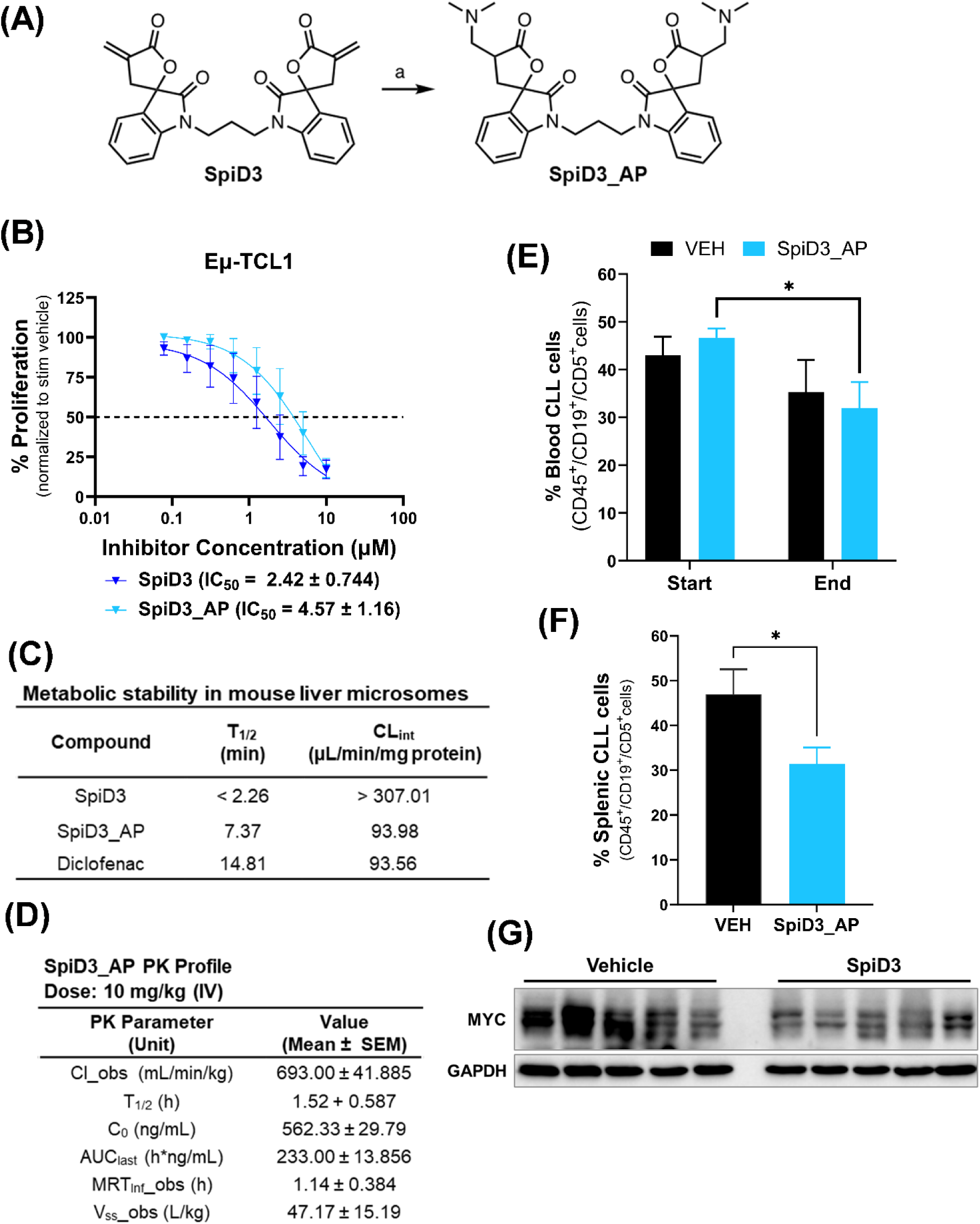
SpiD3 prodrug (SpiD3_AP) displays anti-leukemic activity in Eμ-TCL1 mice. **(A)** Synthesis of SpiD3_AP. Reagents and conditions (a) dimethyl amine (2M in MeOH), MeOH: DCM (2:1), 0°C. **(B)** Spleen-derived malignant B-cells from terminally diseased Eμ-TCL1 mice (n = 7) were stimulated ex vivo with 1X PMA/Ionomycin and treated with increasing concentrations of SpiD3 or SpiD3_AP for 48 h. Proliferation was assessed via MTS assay and normalized to the stimulated vehicle. Error bars and IC_50_ values are shown as mean ± SEM. **(C)** In vitro microsomal stability studies comparing the stability of SpiD3 and SpiD3_AP. Diclofenac (2 µM) was used as a positive control in the metabolic stability study. T_1/2_: half-life, CL_int_: intrinsic clearance. **(D)** Pharmacokinetic (PK) profile of SpiD3_AP administered intravenously (IV) at 10 mg/kg body weight. PK parameters include Cl_obs: clearance observed, T_1/2_: half-life, C_0_: initial concentration, AUC_last_: area under the curve last, MRT_Inf__obs: mean residence time observed, V_ss__obs: steady state volume of distribution. Results are represented as mean ± SEM (n = 3 mice). **(E-G):** Diseased Eμ-TCL1 mice (median age = 10.2 mo) were randomized to receive 10 mg/kg SpiD3_AP or vehicle equivalent (VEH) via IV injection for 3 consecutive days. Equal numbers of male and female mice were used per treatment arm (n = 6 mice/arm). At study end (∼3 h after the last IV injection), mice were sacrificed for tissue harvest. Flow cytometry evaluation of disease burden in blood **(E)** and spleen **(F)**. Error bars are shown as mean ± SEM. **(G)** Representative immunoblot analysis of MYC expression in spleen-derived malignant B-cells from the treated mice. GAPDH served as the loading control. Asterisks indicate significance *vs.* VEH. \**P* < 0.05.

## DISCUSSION

In this study, we introduced a novel spirocyclic dimer, SpiD3, as a potential therapeutic agent for aggressive and indolent B-cell malignancies, including ibrutinib-resistant CLL. We demonstrated SpiD3’s anti-tumor effects independent of TME signals and characterized its unique MoA in CLL cells. Notably, SpiD3 treatment inhibited NF-κB activation and exacerbated unfolded cellular protein loads thereby activating the UPR, culminating in the arrest of protein synthesis and robust CLL cytotoxicity.

Impairment of proper protein folding within the ER results in an accumulation of misfolded or unfolded proteins which are highly cytotoxic [43]. In response, cells activate the UPR to relieve the ER stress induced by increased unfolded/misfolded proteins. However, the inability to do so results in the induction of CHOP protein expression, triggering apoptosis through decreased BCL2 transcription [44]. Leukemic cells have higher rates of protein synthesis [4] that results in higher basal levels of unfolded proteins compared to healthy lymphocytes [9]. Hence, leukemic cells display a vulnerability that can be exploited, providing a unique therapeutic window for selectively inducing apoptosis through sustained UPR activation [10]. In this study, we provide critical proof-of-concept for this strategy using a novel small molecule, SpiD3. Indeed, other studies have shown that ER stress-inducing agents such as thapsigargin [11], and xanthohumol [12] induced CLL cell apoptosis in vitro. Consistent with our previous findings using SpiD7 in ovarian cancer models [10], SpiD3 treatment activated sustained UPR signaling in CLL cells evidenced by an accumulation of unfolded proteins, PERK activation, and increased expression of ATF4, CHOP, and spliced XBP1.

As SEC residues protect against ROS generation [45], the covalent binding of SpiD3 to SEC residues could be a possible mechanism contributing to the elevated oxidative stress and ROS levels observed in our study. Furthermore, accumulation of ROS due to oncogenic (e.g., MYC) activation, the hypoxic TME, and aberrant metabolism induces ER stress leading to mitochondrial dysfunction and apoptosis [43, 44]. Increased ROS levels in CLL cells may sensitize them to agents that further increase oxidative stress [46, 47]. For instance, the pro-apoptotic effects of arsenic trioxide [47] and auranofin [46] were enhanced by HMOX1 induction in CLL cells. We show SpiD3 dramatically increased HMOX1 expression and ROS production in CLL cells. This was accompanied by increased γH2AX expression and cell cycle arrest. Similarly, Lampiasi et al demonstrated that the covalent NF-κB inhibitor, DHMEQ, caused apoptosis in liver cancer cells through the oxidative stress induction and subsequent ROS-mediated DNA damage as increased γH2AX levels were also detected [48]. SpiD3-induced oxidative stress may also contribute to higher levels of ER stress, thus amplifying the pro-apoptotic effects of SpiD3 in CLL cells.

A growing body of evidence suggests that cancer cells’ reliance on heightened protein synthesis stems from irregular translational regulation partially by mTORC1 phosphorylation of 4E-BP1 [32, 34]. Within protective tumor niches, CLL cells display heightened activation of BCR, TLR, NF-κB, E2F and MYC signaling to sustain survival and proliferation [49]. Moreover, studies have reported increased protein synthesis and eIF4F cap-complex formation following BCR stimulation of CLL cells using anti-IgM [5]. Furthermore, oncogenic mRNA processing and translation have emerged as key factors driving CLL disease [4]. Our results demonstrated that SpiD3 effectively decreased MYC expression, induced cell cycle arrest, inhibited protein translation, and modulated NF-κB expression independent of TME stimuli. Overall, these data support SpiD3’s use against highly proliferative and TME-protected CLL cells.

Front-line BTK inhibitors like ibrutinib indirectly inhibit NF-κB activation through blocking upstream BCR signaling; however, direct targeting of NF-κB in CLL has been challenging. In contrast to other NF-κB inhibitors (e.g., TPCA-1), SpiD3 spares healthy lymphocytes and attenuates NF-κB activity in CLL cells, possibly through cross-linking both P65 and IKKβ [14], demonstrating a potential therapeutic window for SpiD3 in CLL. Emerging evidence has strongly implicated NF-κB activation in ibrutinib-resistant CLL [3], emphasizing the importance of therapeutics which directly target NF-κB proteins for relapsed and/or refractory patients. Here, we demonstrated the combination of SpiD3 with ibrutinib was synergistic, suggesting SpiD3 renders CLL cells more sensitive to ibrutinib. Furthermore, SpiD3 exhibited anti-leukemic properties in IR-HG3 cells by impairing survival and proliferative signals (e.g., PRAS, ERK, MYC), bypassing key facets of ibrutinib resistance [39]. Further studies to explore this novel therapeutic modality in other drug resistant CLL models are needed.

To assess SpiD3’s therapeutic capability, we employed the Eµ-TCL1 mouse model for CLL [29] that reliably captures characteristics of aggressive, treatment-resistant human CLL (unmutated *IGVH* disease) and closely recapitulates human CLL in regard to elevated basal levels of ER stress [50] and response to treatment [26]. Despite the brief treatment period, diseased mice treated with SpiD3_AP showed notable reductions in the leukemic burden in both the blood and spleen. This anti-leukemic effect correlated with reduced MYC protein expression suggesting SpiD3_AP could directly attenuate leukemic cell survival. To achieve disease eradication, it’s crucial to develop more durable analogs of SpiD3_AP for prolonged treatments and validate the MoA of SpiD compounds.

Altogether, our data establishes the preclinical activity of a novel spirocyclic dimer, SpiD3, in CLL. SpiD3-mediated cross-linking of cellular proteins targets multiple tumorigenic mechanisms including the UPR and protein translation, ultimately subduing CLL cell survival and proliferation. Furthermore, SpiD3 attenuated NF-κB activation independent of TME support and displayed anti-tumor properties in ibrutinib-resistant cells, underlining its activity against proliferative and/or drug-resistant CLL cells. Concordantly, SpiD3_AP exhibits anti-tumor activity in the Eμ-TCL1 mouse model, warranting further in vivo investigation to characterize the MoA of SpiDs. Collectively, this study strongly supports the development of SpiD3 or SpiD prodrugs as a novel therapeutic approach for aggressive and/or relapsed/refractory B-cell malignancies.

## DATA AVAILABILITY

RNA-sequencing data are deposited at GSE236239. Mass spectrometry data are available via ProteomeXchange with identifier PXD043717 for the click-chemistry experiment and with identifier PXD043688 for the proteomic experiment.

## ACKNOWLEDGMENTS

We are grateful to the patients and the healthy volunteers who provided blood for the above studies. The collection of CLL patient specimens and data used in this study was supported by the integrated Cancer Repository for Cancer Research (iCaRe^2^), developed, and maintained by the Cancer Research Informatics Office and the Clinical Trials Office at the Fred & Panela Buffet Cancer Center (FPBCC). We also thank the Department of Pharmacology and Experimental Neuroscience Elutriation Core at the University of Nebraska Medical Center (UNMC) for the isolation of healthy donor PBMCs used in this study. We also thank Carlo M Croce for access to the Eµ-TCL1 mice. The authors would like to thank Hannah M King, Lelisse T Umeta, and Dalia Y Moore for their technical assistance and Michelle A Lum for cap-binding assay training.

## AUTHORSHIP CONTRIBUTIONS

A.P.E., A.L.S., S.A.S., E.S., and D.E. performed experiments, and analyzed the data; A.P.E., A.N., and D.E. contributed to the conceptualization, experimental design, and interpretation of the results. S.R., S.S., S.K., J.R.M., and A.N. synthesized analog 19, SpiD3, SpiD7, the alkyne-tagged analog 19, the TPE-NMI compound and SpiD3_AP. A.Ka., and M.J.R. assisted with the initial RNA-sequencing analysis and data deposit. A.A.D. assisted with the differential analysis and visualization of the RNA-sequencing and proteomics data. C.R.D., M.A.L, R.G.B., J.M.V. accrued patients, provided clinical insight that helped shape the discussion of the research, and reviewed the manuscript; A.P.E. and D.E. wrote the manuscript. All other authors reviewed and edited the manuscript. D.E. managed and supervised all study aspects. All authors have read and agreed to the published version of the manuscript.

