## Supplemental File for "Novel spirocyclic dimer, SpiD3, targets chronic lymphocytic leukemia survival pathways with potent preclinical effects"

### SUPPLEMENTARY FILE

#### SUPPLEMENTARY TABLES

##### Table S1: Characteristics of the patient-derived CLL samples used in the study

| Patient # | IGHV status | Gender | Age | Treatment | FISH Cytogenetics | | | | | Karyotype* | Figure(s) |
|---|---|---|---|---|---|---|---|---|---|---|---|
| | | | | | Deletion (17p) | Deletion (13q) | Deletion (11q) | Deletion (6q) | Trisomy 12 | | |
| 1 | Unmut | M | 47 | Naïve | Neg | Neg | Neg | Neg | Neg | Normal | 5B |
| 2 | Unmut | M | 55 | Naïve | Neg | Neg | Pos | Pos | Neg | Complex | 5B |
| 3 | Unmut | F | 76 | Naïve | Neg | Neg | Pos | Neg | Neg | Normal | 5A-B |
| 4 | Unmut | F | 65 | Naïve | Neg | Neg | Neg | Neg | Pos | Normal | 5B |
| 5 | NA | M | 81 | Treated | Neg | Neg | Neg | Neg | Neg | Normal | 5B |
| 6 | NA | F | 68 | Treated | Neg | Neg | Neg | Neg | Pos | Normal | 5A |
| 7 | Mut | M | 94 | Treated | Neg | Neg | Neg | Neg | Pos | Normal | 5A |
| 8 | Unmut | M | 76 | Treated | Pos | Neg | Neg | Neg | Neg | Complex | 5A |
| 9 | Mut | M | 47 | Naïve | Neg | Neg | Neg | Neg | Pos | Complex | 5A |
| 10 | Unmut | F | 62 | Naïve | Neg | Neg | Neg | Neg | Pos | Normal | 5A |
| 11 | Unmut | M | 76 | Treated | Neg | Neg | Neg | Neg | Pos | Normal | 5A-B |
| 12 | Unmut | M | 45 | Naïve | Neg | Neg | Pos | Neg | Pos | Normal | 5A |
| 13 | NA | M | 57 | Naïve | Pos | Neg | Neg | Neg | Neg | Complex | 5A |
| 14 | Unmut | M | 70 | Naïve | Neg | Neg | Neg | Neg | Neg | Normal | 5A |
| 15 | Unmut | M | 34 | Naïve | Neg | Neg | Neg | Neg | Pos | Normal | 5A |
| 16 | Unmut | F | 60 | Treated | Neg | Neg | Pos | Neg | Neg | Normal | 5A |
| 17 | Unmut | M | 47 | Naïve | Neg | Neg | Neg | Neg | Neg | Normal | 5A |
| 18 | Unmut | F | 52 | Naïve | Neg | Neg | Neg | Neg | Neg | Complex | 5A |
| 19 | Mut | F | 66 | Treated | Neg | Pos | Neg | Neg | Neg | Normal | 5B |
| 20 | Unmut | F | 68 | Treated | Neg | Pos | Pos | Neg | Neg | NA | 5B |
| 21 | Mut | F | 78 | Treated | Neg | Pos | Pos | Neg | Neg | Complex | 5B |
| 22 | Mut | M | 80 | Naïve | Neg | Pos | Neg | Neg | Neg | NA | 5C |
| 23 | Unmut | F | 53 | Naïve | Neg | Pos | Neg | Neg | Neg | Normal | 5B-D |
| 24 | NA | M | 66 | Naïve | NA | NA | NA | NA | NA | NA | 5D |
| 25 | Mut | M | 78 | Naïve | Neg | Pos | Neg | Neg | Neg | Normal | 5B |
| 26 | Unmut | M | 65 | Naïve | Neg | Neg | Neg | Neg | Neg | NA | 5B, 5D |
| 27 | Unmut | F | 65 | Naïve | Neg | Pos | Pos | Neg | Neg | NA | 5B-D |
| 28 | Mut | M | 45 | Treated | Neg | Neg | Neg | Neg | Neg | NA | 5D |
| 29 | NA | M | 66 | Naïve | Neg | Neg | Neg | Neg | Neg | Normal | 5C, 5E |
| 30 | NA | F | 58 | Naïve | NA | NA | NA | NA | NA | NA | 5B-C |
| 31 | Unmut | M | 45 | Treated | Neg | Pos | Neg | Neg | Neg | Normal | 5B-C |
| 32 | Unmut | M | 72 | Naïve | Neg | Neg | Neg | Neg | Neg | Normal | 5D |
| 33 | Unmut | M | 70 | Naïve | Pos | Neg | Neg | Neg | Neg | NA | 5B, 5E |
| 34 | Unmut | F | 68 | Treated | Neg | Neg | Neg | Neg | Neg | Normal | 5A, 5C-D |
| 35 | Unmut | M | 68 | Naïve | Neg | Neg | Neg | Pos | Neg | NA | 5B |
| 36 | NA | F | 75 | Treated | Pos | Pos | Neg | Neg | Neg | Complex | 5D |
| 37 | Mut | M | 50 | Naïve | Neg | Pos | Neg | Neg | Neg | Normal | 5B |
| 38 | Unmut | M | 61 | Treated | Neg | Neg | Neg | Neg | Pos | Normal | 5D |
| 39 | NA | F | 82 | Treated | Neg | Pos | Neg | Neg | Neg | Normal | 5B |
| 40 | Unmut | M | 59 | Naïve | Neg | Pos | Pos | Neg | Neg | NA | 5B |

Unmut: unmutated *IGHV*, Mut: mutated *IGHV*; M: male, F: female; Neg: negative, Pos: positive; NA: not available. *Complex karyotype is defined as >3 cytogenetic abnormalities

**Table S2: List of primary antibodies used for immunoblotting assays in the study**

| Primary Antibodies | Catalog |
|---|---|
| **Cell Signaling Technology** | |
| p4E-BP1 (Ser65) | 9456 |
| 4E-BP1 | 9644 |
| ATF4 | 11815 |
| p-BTK (Tyr223) | 87457 |
| BTK | 8547 |
| CHOP | 2895 |
| p-eIF2α (Ser51) | 3398 |
| eIF2α | 5324 |
| eIF4A1 | 2490 |
| eIF4E | 2067 |
| eIF4G1 | 2469 |
| p-ERK1/2 (Thr202/Tyr204) | 4377 |
| ERK1/2 | 4695 |
| GAPDH | 5174 |
| γH2A.X (Ser139) | 9718 |
| HO-1/HMOX1 | 43966 |
| IKKβ | 8943 |
| IRE1α | 3294 |
| MCL1 | 5453 |
| MYC | 5605 |
| P21 | 2947 |
| P52 | 4882 |
| PARP | 9542 |
| PDCD4 | 9535 |
| PERK | 5683 |
| p-PRAS (Thr246) | 13175 |
| PRAS | 2691 |
| RELB | 10544 |
| **Invitrogen** | |
| LAMIN B1 | 702972 |
| XBP1 (spliced/unspliced) | PA5-27650 |
| **Santa Cruz Biotechnology** | |
| β-ACTIN | sc-47778 |
| IKKα | sc-52932 |
| P65 | sc-8008 |
| **Sigma** | |
| α-TUBULIN | T5201 |

#### SUPPLEMENTARY METHODS

##### Click chemistry

OSU-CLL cells (2e$^6$ cells/mL) were treated with the alkyne-tagged derivative of analog 19 (10 μM) for 2 h at 37°C. Lysates were prepared in lysis buffer [150 mM NaCl, 50 mM HEPES pH 7.4, 1 % Igepal CA-630 and 1% sodium dodecyl sulfate (SDS)] that contained fresh protease and phosphatase inhibitors (1:100; Sigma-Aldrich; St. Louis, MO), and centrifuged at 13,000 rpm for 10 min at 4°C. Protein concentrations of supernatants were determined by the BCA assay

(ThermoFisher Scientific; Waltham, MA) and 100 µg of protein was reacted with 10 µL of BTTAA ligand (40 mM), 10 µL of Copper (II) Sulfate + Protectant, 10 µL of reducing agent (20 mg) and 10 µL of 5 mM TAMRA biotin azide (Click Chemistry Tools; Scottsdale, AZ). The mixture was rotated for 90 min at room temperature and then the proteins were precipitated by adding equal volumes of 3:1 chloroform/methanol and the pellets were washed with ice-cold methanol. The pellets were then resuspended in resuspension buffer (150 mM NaCl, 50 mM Tris, and 1% SDS) and incubated with streptavidin agarose resin (Click Chemistry Tools) for 2 h on a rotator. The resin was centrifuged and washed with resuspension buffer, then 1 % SDS in phosphate-buffered saline (PBS), and then with only PBS.

For the mass spectrometry (MS) analysis, the resin was resuspended in 500 µL of DTT (10 mM) and heated at 70°C for 15 min. Following a 5 min centrifugation, 1 mL of iodoacetamide solution (40 mM) was added and incubated in the dark for 30 min. The resin was pelleted and washed with PBS and then added to 300 µL of digestion buffer (2 mM of $CaCl_2$, 100 mM Tris, and 0.5 M Urea) and 3 µg of trypsin and incubated overnight on a rotator. The next day, the resin was washed with PBS, and supernatant was collected and concentrated using a SpeedVac. The peptides were cleaned from salts and detergents and analyzed using a high-resolution mass spectrometry nano-LC-MS/MS Tribrid system, Orbitrap Fusion™ Lumos™ coupled with UltiMate 3000 HPLC system (ThermoFisher Scientific) at the UNMC Proteomics Core. Approximately 1 µg of peptides were run on the pre-column (Acclaim PepMap™ 100, 75µm × 2cm, nanoViper; ThermoFisher Scientific) and the analytical column (Acclaim PepMap™ RSCL, 75 µm × 50 cm, nanoViper; ThermoFisher Scientific). The samples were eluted using a 155-min linear gradient of ACN (4-45 %) in 0.1 % FA. All MS/MS samples were analyzed using Proteome Discoverer (ThermoFisher Scientific, vs 2.2.). Sequest HT was set up to search the SwissProt database (selected for Human, 2021_04, 20395 entries) assuming the digestion enzyme trypsin. The parameters for Sequest HT were set as follows: Enzyme: trypsin, Max missed cleavage: 2, Precursor mass tolerance: 10 ppm, Peptide tolerance: ± 0.6 Da, Fixed modifications: carbamidomethyl (C); Dynamic modifications: oxidation (M). Consensus workflow was chosen for enhanced annotation LFQ and precursor quantitation. The parameters for Precursor ion quantifier were set as follows: peptides to use: unique + razor; precursor abundance: intensity; normalization mode: total peptide amount; scaling mode: on all average; peptide confidence: high; target *FDR* (strict): 0.01: target *FDR* (relaxed): 0.05. The data was analyzed by comparing the list of proteins from each experiment in a Venn diagram and the list of proteins that were found in at least two out of the three biological replicates/samples were put into EnrichR [1-3] for pathway analysis. The pathways were graphed in a bubble plot based on their -$Log_{10}$ *P* values. MS data is available via ProteomeXchange with identifier PXD043717.

For the immunoblotting, the resin was resuspended in 500 µL of regeneration buffer (0.1 M HCL glycine, pH 2.8) and incubated for 10 min at room temperature. The resin was centrifuged, and the eluents were collected and concentrated using an Amicon 10 kDa molecular weight cutoff filter (Sigma-Aldrich). The concentrated samples were evaporated to dryness in a SpeedVac for 10 h at 4°C. The lyophilized samples were dissolved in PBS (25 µL) and 6X sample loading dye was added (5 µL) and vortexed. The input lysate was dissolved in 15 µL of PBS and 2X loading dye was added (15 µL, Bio-Rad; Hercules, CA) and vortexed. All samples were heated to 90°C for 5 min, subjected to SDS-PAGE, and subsequent immunoblotting.

**Mass spectrometry (MS)**

OSU-CLL ($2e^6$ cells/mL) were treated for 24 h with 1 µM SpiD3 or vehicle equivalent (DMSO; n = 3 replicates), and whole cell lysates were extracted and quantified according to the immunoblotting protocol. Using the tandem mass tag (TMT) 10-plex Mass Tag Labeling Kits (ThermoFisher Scientific), 100 µg of each sample was diluted with 100 mM TEAB and labeled. Pierce Quantitative Colorimetric Peptide Assay (ThermoFisher Scientific) was used for peptide quantification and the labeled peptides were run on the Orbitrap Fusion™ Lumos™ (ThermoFisher Scientific) mass spectrometry machine at the UNMC Proteomics Core and then analyzed using Proteome Discoverer (ThermoFisher Scientific, vs 2.2). Sequest HT was set up to search the SwissProt database (selected for Human, 2021_04, 20395 entries) assuming the digestion enzyme trypsin. The parameters for Sequest HT were set as follows: Enzyme: trypsin; Max missed cleavage: 2; Precursor mass tolerance: 10 ppm; Peptide tolerance: ± 0.6 Da; Fixed modifications: carbamidomethyl (C); Dynamic modifications: oxidation (M). Consensus workflow was chosen for enhanced annotation LFQ and precursor quantitation. The parameters for Precursor ion quantifier were set as follows: peptides to use: unique + razor; precursor abundance: intensity; normalization mode: total peptide amount; scaling mode: on all average; peptide confidence: high; target *FDR* (strict): 0.01; target *FDR* (relaxed): 0.05. Differential protein expression analysis was performed using the limma package [4]. Functional annotation and pathway enrichment analysis for the differentially expressed proteins ($P > 0.05$) were evaluated using MSigDB [5-7]. MS data is available via ProteomeXchange with identifier PXD043688.

**Weighted gene co-expression network analysis (WGCNA)**

Using the top 75% most variably expressed genes (n = 15,005), a pairwise gene correlation matrix was calculated with a Pearson correlation analysis, which was transformed into a signed weighted matrix to produce an adjacency matrix after raising values by an exponent beta (β = 10). Then the adjacency was transformed into a topological overlap matrix (TOM). The dynamic tree cut method was used for module identification from the hierarchical clustering of genes using 1-TOM as the distance measure with a deepSplit value of 2 and a minimum size cutoff of 15 genes. Finally, modules and their relationship to drug concentration (1 μM or 2 μM SpiD3) or vehicle (DMSO) were identified using Pearson correlation analysis between the modules and external traits. Functional annotation of identified modules was performed using tools provided by the WGCNA package [8].

**Chemotaxis assay**

Following 1 h pretreatment with DMSO, SpiD3, or ibrutinib, OSU-CLL (25,000 cell/mL) cells were placed on 5 μM trans-well inserts in 24-well plates containing 200 ng/mL CXCL-12 (PeproTech; Cranbury, NJ). After 6 h, the number of cells that migrated through the insert towards the chemokines were counted by flow cytometry analysis. The chemotaxis index represents the number of cells that migrated toward the indicated chemokine divided by the number of cells that migrated with no chemokine present for each treatment condition.

**Real time quantitative PCR**

RNA was extracted from CLL cells after inhibitor treatment using the miRNeasy Mini Kit (Qiagen; Hilden, Germany). RNA (1 mg) was used to make cDNA using iScript cDNA Synthesis Kit (Bio-Rad) following the manufacture's protocol. mRNAs were quantified with iTaq Universal SYBR Green Supermix (Bio-Rad) on the QuantStudio®3 Real-Time PCR System (Applied Biosystems, Waltham, MA). Target gene expression was determined using the $2^{-\Delta\Delta CT}$ method [9] and presented relative to *GAPDH*. Primers were as follows: *IRE1* forward: 5'AGACTTTGTCATCGGCCTTTGCAG3', reverse: 5'-ATTCACTGTCCACAGTCACCACCA-3'; *EIF2AK3 (PERK)* forward: 5'-GCAACAACGTTTATTGTGCGCAGG-3', reverse: 5'-AAACAACTCCAAAGCCACCACGTC-3'; *ATF4* forward: 5'-AAGCCTAGGTCTCTTAGATG-3', reverse: 5'-TTCCAGGTCATCTATACCCA-3'; *DDIT3 (CHOP)* forward: 5'-TCTTCACCACTCTTGACCCTGCTT-3', reverse: 5'-GTTCTTTCTCCTTCATGCGCTGCT-3'; and *GAPDH* forward: 5'-TGAAGGTCGGAGTCAACGGA-3', reverse: 5'-CCATTGATGACAAGCTTCCCG-3'.

**Immunoblot assays**

The NE-PER™ extraction kit (ThermoFisher Scientific) was used for nuclear and cytoplasmic lysate preparation following manufacturer instructions. For the cap-binding assay, whole cell lysate (WCL) was incubated with agarose-immobilized $m^7$GTP cap analogs (Jena Bioscience; Germany) to capture eIF4E and its binding partners (4E-BP1, eIF4G) as previously described [10]. The WCL was prepared using protein lysis buffer (20 mM Tris pH 7.4, 150 mM NaCl, 1% Igepal CA-630, 5 mM EDTA) containing protease and phosphatase inhibitor cocktails and phenylmethyl sulfonyl fluoride (Sigma-Aldrich).

BCA protein analysis (ThermoFisher Scientific) was used to determine equal concentrations of protein for each sample lysate (whole cell, nuclear, or cytoplasmic). All samples were heated to 90°C for 5 min, subjected to SDS-PAGE, and subsequent immunoblotting.

**SpiD3 prodrug (SpiD3_AP) synthesis details**

**General methods - Chemistry:** All reagents were purchased from commercial sources and used without further purification. Flash chromatography was carried out on silica gel (200–400 mesh). Thin-layer chromatography was run on pre-coated ANALTECH plates and observed under UV light at 254 nm. Column chromatography was performed with silica gel (230-400 mesh, grade 60, Fisher Scientific; Hampton, NH). $^1$H NMR (400 MHz) and $^{13}$C NMR (100 MHz) spectra were recorded in chloroform-d or DMSO-*d*6 on a Bruker-400 spectrometer (DMSO-*d*6 was 2.50 ppm for $^1$H and 39.55 ppm for $^{13}$C, and $CDCl_3$ was 7.26 ppm for $^1$H and 77.23 ppm for $^{13}$C). Proton and carbon chemical shifts were reported in ppm relative to the signal from residual solvent proton and carbon. The data are presented as follows: chemical shift, multiplicity (s = singlet, d = doublet, t = triplet, q = quartet, p = pentet, m = multiplet and/or multiple resonances), coupling constant in hertz (Hz), and integration. The purified compounds were further confirmed by high-resolution mass spectrometry (HRMS) analysis using the Agilent 6230 time-of-flight LC/MS (LC/TOF) system (Santa Clara, CA).

**1',1'''-(propane-1,3-diyl)bis(4-((dimethylamino)methyl)-3,4-dihydro-5H-spiro[furan-2,3'-indoline]-2',5-dione) (SpiD3_AP):** Dichloromethane (4 mL), MeOH (10 mL) was added to a stirred solution of SpiD3 (250 mg, 0.53 mmol). The mixture was stirred at 0°C for 10 min, followed by the addition of a 2M solution of dimethylamine (0.611 mL, 1.22 mmol). The reaction was allowed to stir for 5 min at 0°C. The solvent and excess dimethylamine were removed under vacuum. The crude product was further dissolved in 1 mL of tetrahydrofuran, to which *n*-pentane was slowly added until white precipitates formed. The resulting solid was filtered, washed with *n*-pentane, and dried; Yield = 84.5% (251 mg);$^1$H NMR (400 MHz, $CDCl_3$) δ (mixture of diastereomers) 7.37 (dt, $J$ = 13.5, 7.2 Hz, 4H), 7.14 (dd, $J$ = 15.0, 7.4 Hz, 2H), 6.93 – 6.72 (m, 2H), 4.00 – 3.66 (m, 4H), 3.64 – 3.50 (m, 1H), 3.30 – 3.14 (m, 1H), 2.88 (m, $J$ = 7.7, 4.5 Hz, 2H), 2.81 – 2.52 (m, 4H), 2.49 – 2.35 (m, 2H), 2.31 (dd, $J$ = 4.0, 2.0 Hz, 12H), 2.14 (m, 2H); $^{13}C$ NMR (100 MHz, $CDCl_3$) δ 177.3, 177.2, 176.9, 174.6, 174.5, 173.9, 142.9, 142.8, 142.3, 131.3, 131.2, 127.8, 126.2, 124.7, 124.0, 123.7, 109.0, 108.9, 81.1, 80.8, 80.7, 60.7, 59.9, 45.6, 45.4, 39.1, 38.6, 38.0., 37.9, 37.8, 37.6, 37.5, 36.9, 35.6, 25.1, 25.0, 24.8; HRMS(ESI +): m/z calculated for $C_{31}H_{36}N_4O_6$, 561.2708, found, 561.1647 $[M + H]^+$.

**$^1H$ NMR (400 MHz, $CDCl_3$)**

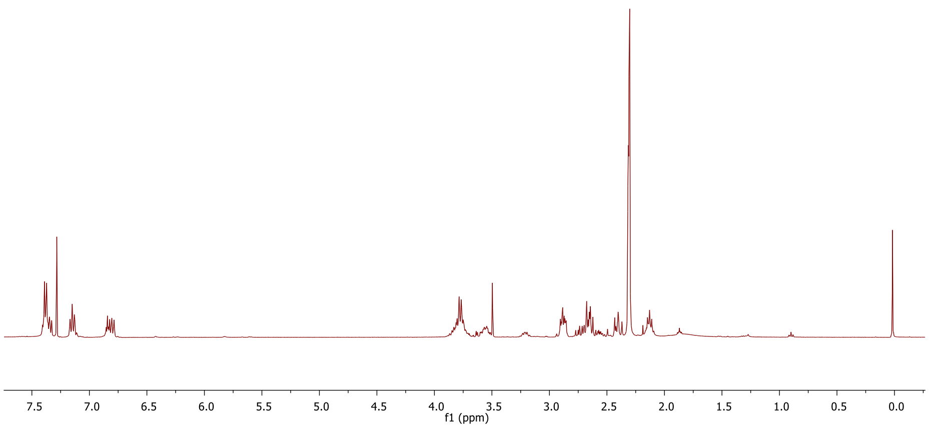


**$^{13}C$ NMR (100 MHz, $CDCl_3$)**

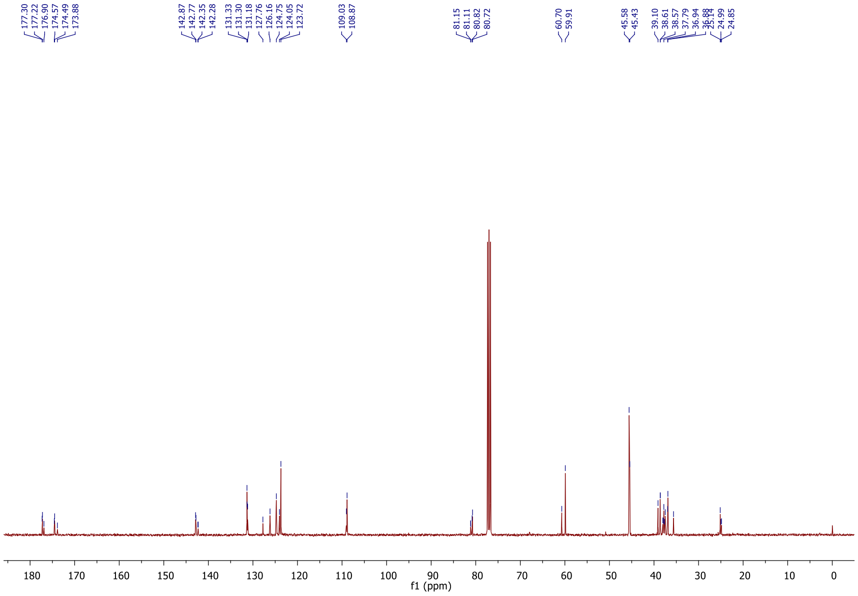

**HRMS**

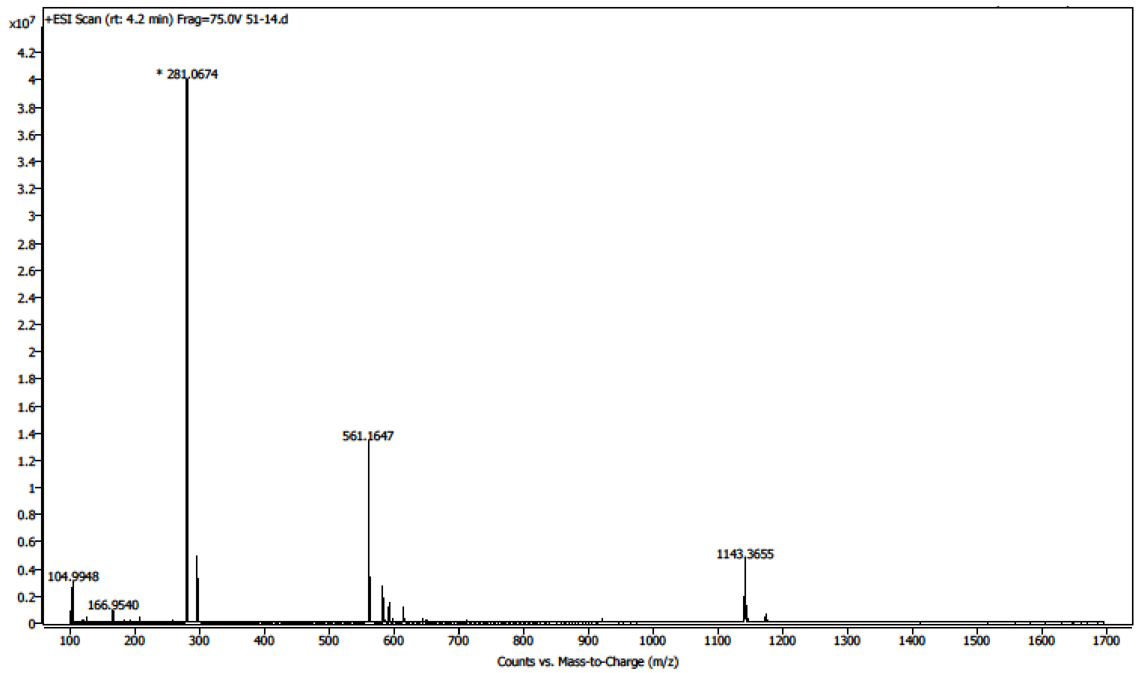


**Metabolic stability studies**

The in vitro metabolic stability of SpiD3 and SpiD3_AP was determined using mouse liver S9 fraction (XenoTech; Kansas City, KS) adapting previously reported methods [11, 12]. Diclofenac was used as a positive control for in vitro metabolic stability study. Sample analysis was performed by liquid chromatography-tandem mass spectrometry (SCIEX QTRAP 4000 LC-MS/MS System).

**Pharmacokinetics (PK) studies**

Intravenous (I.V.) dose pharmacokinetic (PK) studies were performed in CD1 mice. The intravenous dose of SpiD3_AP was 10 mg/kg body weight. Sample analysis was performed by liquid chromatography-tandem mass spectrometry (SCIEX QTRAP 4000 LC-MS/MS System) adapting previously reported methods [11-15]. PK parameters were determined using non-compartmental analyses module of WinNonlin® 1.5.

#### SUPPLEMENTARY DATA FIGURES

**Supplementary Figure S1. RNA-sequencing analysis of SpiD3-treated OSU-CLL cells**

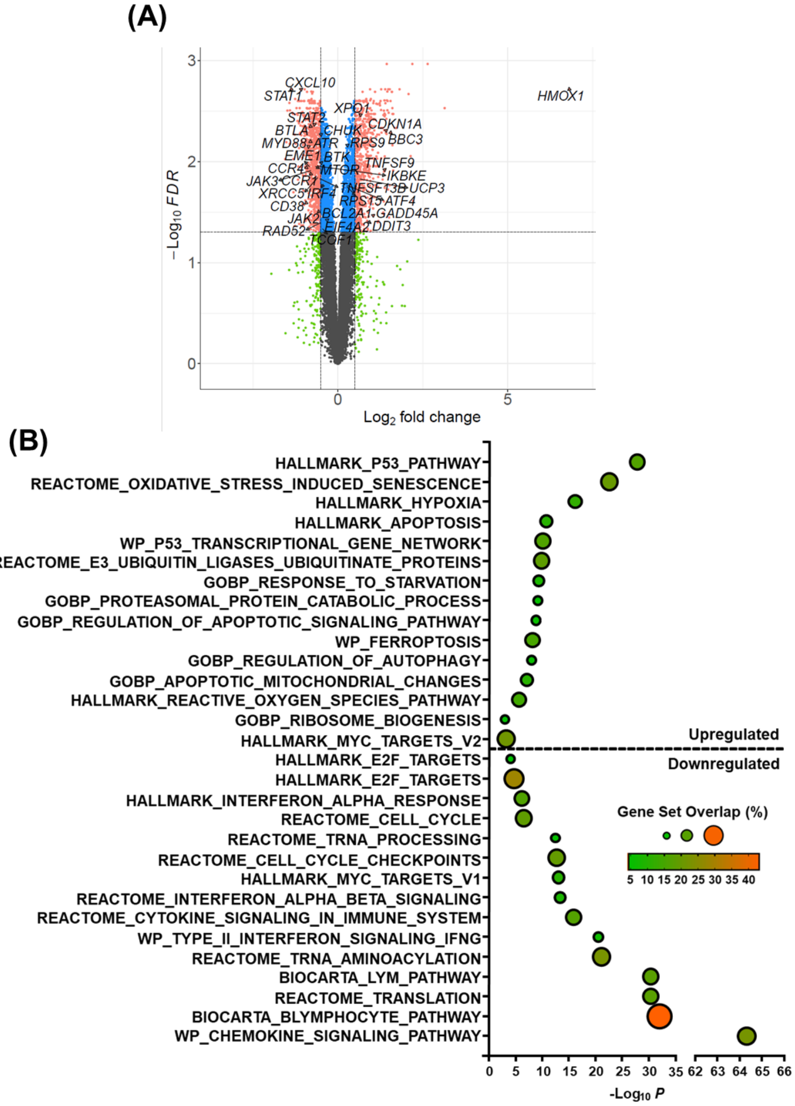


**Supplementary Figure S1. RNA-sequencing analysis of SpiD3-treated OSU-CLL cells. (A)** Volcano plot of differentially expressed genes (DEGs) in SpiD3-treated OSU-CLL cells (1 μM, 4 h) with select disease-relevant genes labeled. Only genes that meet both the statistical significance (*FDR* < 0.05) and fold-change ($|Log_2 FC| > 0.5$) were used for downstream analysis (red). Genes that meet only statistical significance (blue), only fold-change (green), or neither threshold (grey) are shown for comparison. **(B)** Gene set enrichment analysis using the top 500 DEGs (*P* < 0.05) following treatment with 1 μM SpiD3 (4 h, n = 3 independent RNA-seq experiments).

**Supplementary Figure S2. Proteomic analysis of SpiD3-treated CLL cells using TMT-labeling**

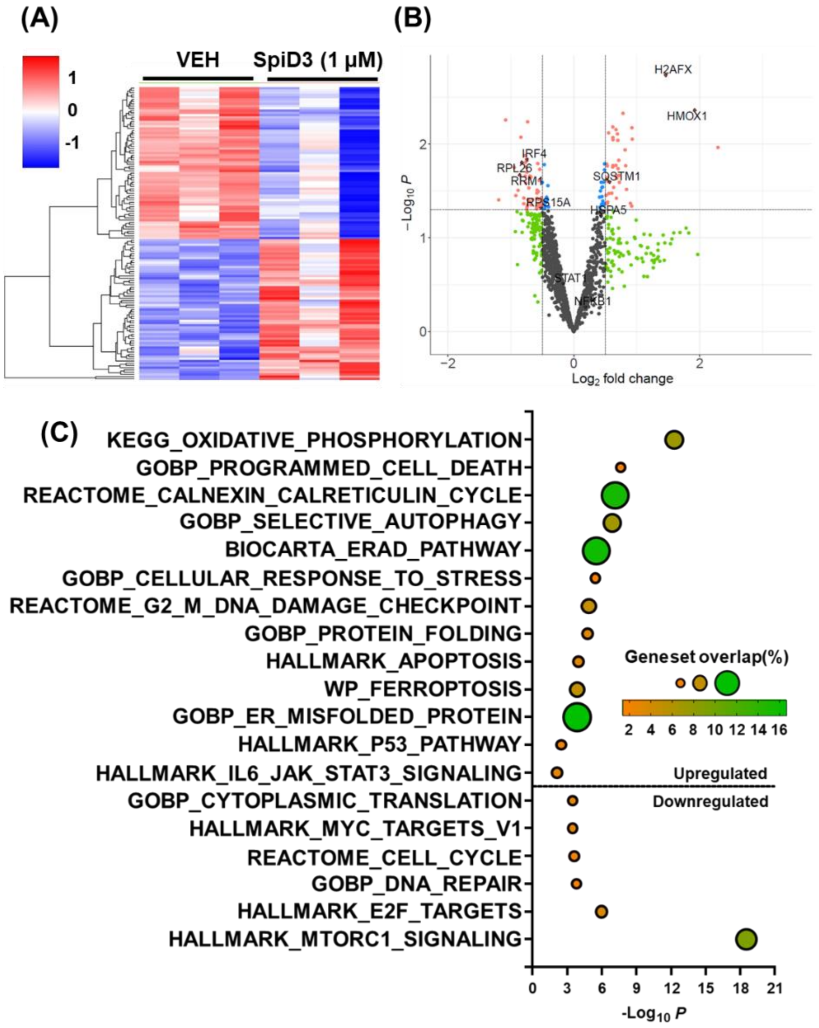


**Supplementary Figure S2. Proteomic analysis of SpiD3-treated CLL cells using TMT-labeling. (A)** Hierarchical clustering of differentially expressed proteins (n = 131, $P$<0.05) in OSU-CLL cells treated with DMSO vehicle (VEH) or SpiD3 (1 μM) for 24 h (n = 3 independent experiments). **(B)** Volcano plot of differentially expressed proteins in SpiD3-treated OSU-CLL cells with select disease-relevant proteins labeled. Only proteins that meet both the statistical significance ($P$ < 0.05) and fold-change ($|\log_2 FC| > 0.5$) were used for downstream analysis (red). Genes that meet only statistical significance (blue), only fold-change (green), or neither threshold (grey) are shown for comparison. **(C)** Gene set enrichment analysis of the differentially expressed proteins following SpiD3 treatment.

**Supplementary Figure S3. SpiD3 inhibits CLL cell chemotaxis and induces transcription of UPR genes**

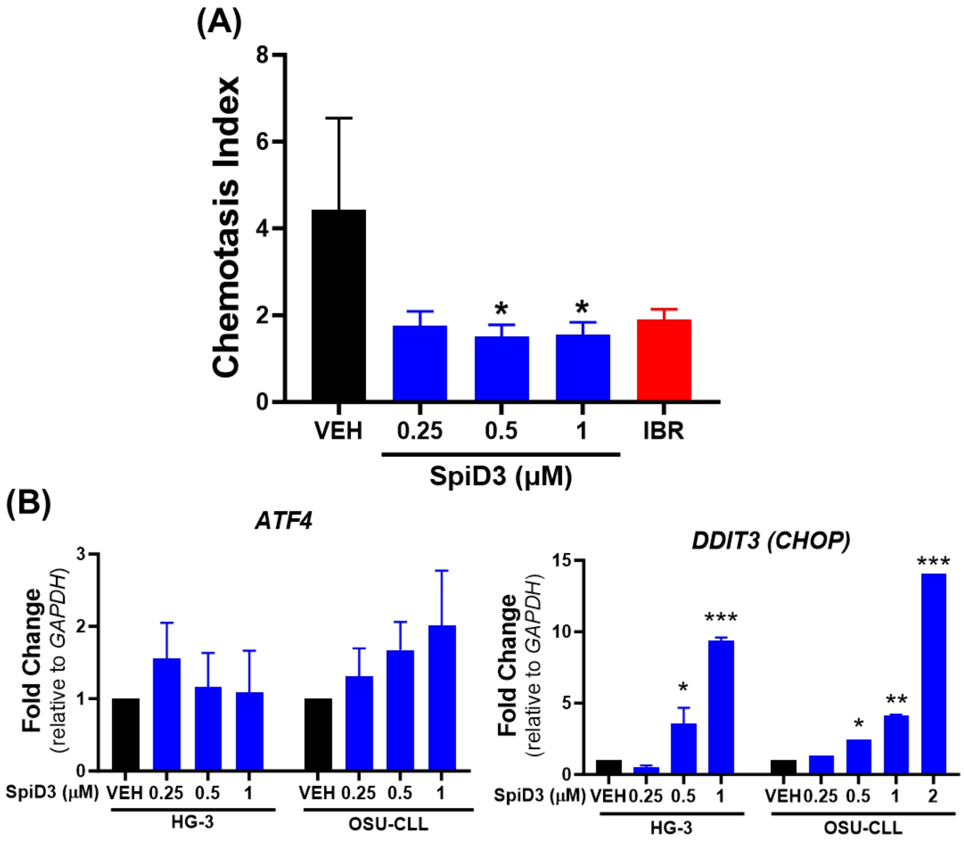


**Supplementary Figure S3. SpiD3 inhibits CLL chemotaxis and induces transcription of UPR genes. (A)** OSU-CLL cells were pre-treated with SpiD3 (0.25-1 µM), ibrutinib (IBR, 1 µM), or equivalent DMSO vehicle (VEH) for 1 h and allowed to migrate for 6 h through trans-well inserts toward 200 ng/mL CXCL-12. The chemotaxis index represents the number of cells that migrated toward the indicated chemokine divided by the number of cells that migrated with no chemokine present for each treatment condition (n = 3 independent experiments). **(B)** Quantitative real-time PCR analysis of *ATF4* and *DDIT3 (CHOP)* in HG-3 and OSU-CLL cells treated with equivalent VEH or SpiD3 (0.25-2 µM) for 4 h (n = 2-3 independent experiments/cell line). Transcript expression is normalized to the housekeeping gene (*GAPDH*). Data are shown as fold change to VEH (mean ± SEM). Asterisks denote significance *vs.* VEH: *$P < 0.05$, **$P < 0.01$, ***$P < 0.001$.

**Supplementary Figure S4: SpiD3 inhibits cap-dependent protein translation in CLL cells**

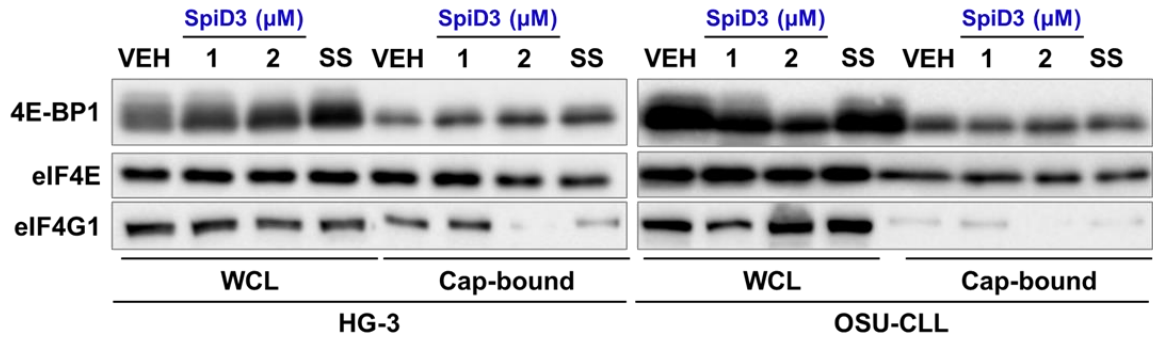


**Supplementary Figure S4: SpiD3 inhibits cap-dependent protein translation in CLL cells.** HG-3 and OSU-CLL cells were treated with VEH, SpiD3 (1, 2 µM), or cultured in serum-deprived media (SS) for 8 h. Cell lysates were incubated with 7-methyl-GTP-Sepharose resin (Cap-bound). Eluted cap-bound proteins and whole cell lysates (WCL) were analyzed via immunoblot for 4E-BP1, eIF4E and eIF4G1 (n = 3 independent experiments).

**Supplementary Figure S5: SpiD3 inhibits NF-κB activity and diminishes CD40L-induced survival signaling in CLL**

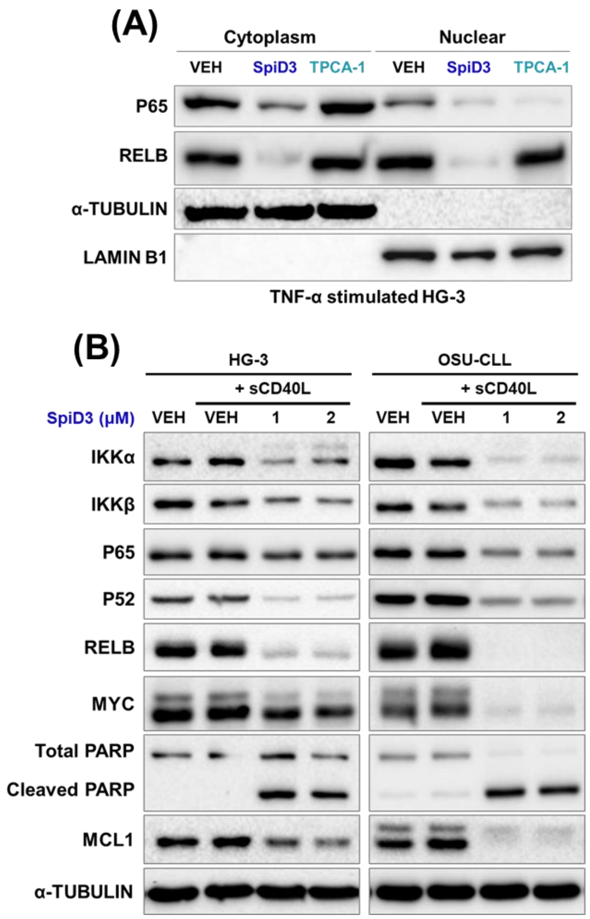


**Supplementary Figure S5: SpiD3 inhibits NF-κB activity and diminishes CD40L-induced survival signaling in CLL.** **(A)** HG-3 cells were treated with SpiD3 (2 μM), TPCA-1 (10 μM), or equivalent DMSO vehicle (VEH) for 4 h and stimulated with TNF-α (20 ng/mL) during the last 15 min of treatment (n = 4 independent experiments). Cytoplasmic and nuclear fractions were subjected to immunoblotting probing for P65 and RELB. α-TUBULIN served as the cytoplasmic fraction loading control and LAMIN B1 served as the nuclear fraction loading control. **(B)** Immunoblot analysis of the indicated proteins in whole cell lysates of HG-3 and OSU-CLL cells treated with SpiD3 (1, 2 μM) for 4 hr and co-currently stimulated with sCD40L (500 ng/mL). α-TUBULIN served as the loading control (n = 4 independent experiments/cell line).

**Supplementary Figure S6: SpiD3 spares healthy stromal and lymphoid cells**

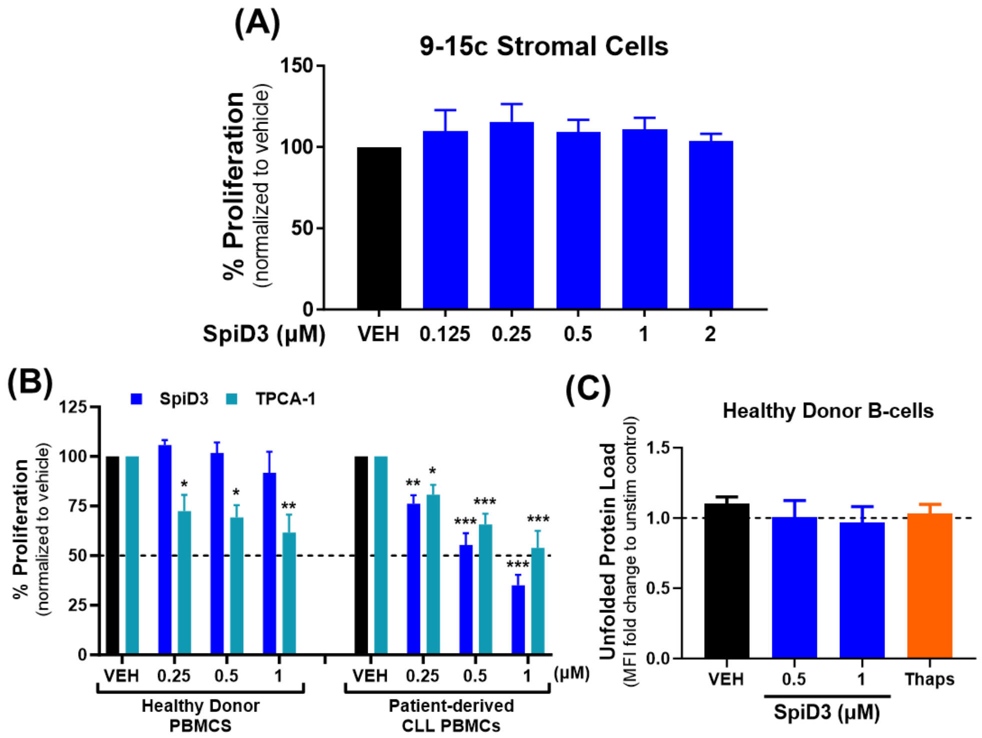


**Supplementary Figure S6: SpiD3 spares healthy stromal and lymphoid cells. (A)** 9-15c bone marrow-derived stroma cells were treated with SpiD3 (48 h). Proliferation was assessed via MTS assay and results are given as % proliferation normalized to vehicle (n = 4 independent experiments). **(B)** Healthy donor PBMCs (n = 5) or patient-derived CLL PBMCs (n = 8; > 92% CLL B-cells) were treated with the indicated amounts of SpiD3 or TPCA-1 with co-current CpG ODN 2006 stimulation (CpG; 3.2 µM) for 48 h. Proliferation was assessed via MTS assay and results are given as % proliferation normalized to the CpG-stimulated control. Data are shown as mean ± SEM. Asterisks denote significance *vs*. VEH: *$P < 0.05$, **$P < 0.01$, ***$P < 0.001$. **(C)** UPR induction in healthy donor B-cells (n = 7) was evaluated following a 24 h ex vivo treatment with SpiD3 or thapsigargin (Thaps; 1 µM) under co-current CpG stimulation (3.2 µM) via incubation with TPE-NMI dye. Data are represented as fold change in TPE-NMI median fluorescence intensity (MFI) compared to the unstimulated control (dashed line).

**Supplementary Figure S7: Studies with SpiD3_AP in CLL**

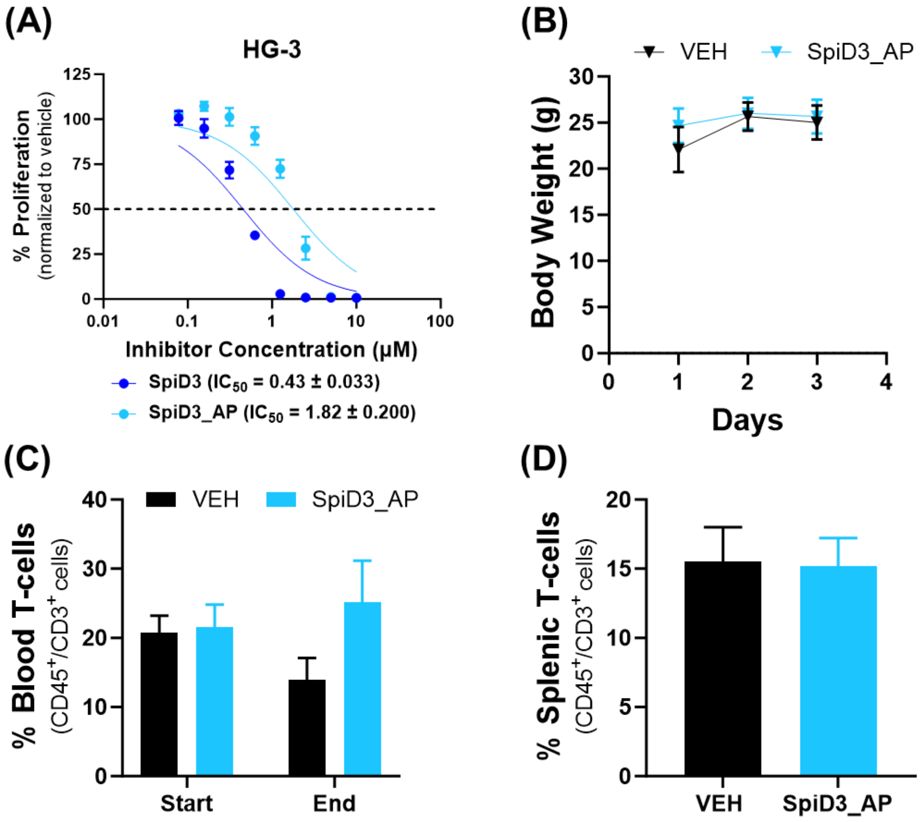


**Supplementary Figure S7: Studies with SpiD3_AP in CLL. (A)** Proliferation of HG-3 CLL cell line was determined by MTS assay following treatment with increasing concentrations of SpiD3 or SpiD3 prodrug (SpiD3_AP) for 72 h (n = 3 independent experiments). Proliferation was assessed via MTS assay and normalized to vehicle. $IC_{50}$ values are shown as mean ± SEM. **(B-D):** Diseased Eμ-TCL1 mice (median age = 10.2 mo) were randomized to receive 10 mg/kg SpiD3_AP or vehicle equivalent (VEH) via intravenously (IV) injection for 3 consecutive days. Equal numbers of male and female mice were used per treatment arm (n = 6 mice/arm). Mouse body weight was monitored throughout the study **(B)**. At study end (~3 h after the last IV injection), mice were sacrificed for tissue harvest. Flow cytometry evaluation of T-cells in blood **(C)** and spleen **(D)**. Data are shown as mean ± SEM.
